# Investigation of gating in outer membrane porins provides new perspectives on antibiotic resistance mechanisms

**DOI:** 10.1101/2021.09.09.459668

**Authors:** Archit Kumar Vasan, Nandan Haloi, Rebecca Joy Ulrich, Mary Elizabeth Metcalf, Po-Chao Wen, William W. Metcalf, Paul J. Hergenrother, Diwakar Shukla, Emad Tajkhorshid

**Affiliations:** NIH Center for Macromolecular Modeling and Bioinformatics, Beckman Institute for Advanced Science and Technology, University of Illinois at Urbana-Champaign, Urbana, IL 61801, USA; Center for Biophysics and Quantitative Biology, University of Illinois at Urbana-Champaign, Urbana, IL 61801, USA; Department of Biochemistry, University of Illinois at Urbana-Champaign, Urbana, IL 61801, USA; Department of Microbiology, University of Illinois at Urbana-Champaign, Urbana, IL 61801, USA; Department of Chemistry, University of Illinois at Urbana-Champaign, Urbana, IL 61801, USA; Carl R. Woese Institute for Genomic Biology, University of Illinois at Urbana-Champaign, Urbana, IL 61801, USA; Cancer Center at Illinois, University of Illinois at Urbana-Champaign, Urbana, IL 61801, USA; Department of Chemical and Biomolecular Engineering, University of Illinois at Urbana-Champaign, Urbana, IL 61801, USA; National Center for Supercomputing Applications, University of Illinois at Urbana-Champaign, Urbana, IL 61801, USA

**Author notes:** These authors contributed equally to this work (Archit Kumar Vasan, Nandan Haloi).

## Abstract

Gram-negative bacteria pose a serious public health concern, primarily due to a higher frequency of antibiotic resistance conferred to them as a result of low permeability of their outer membrane (OM). Antibiotics capable of traversing the OM typically permeate through OM porins; thus, understanding the permeation properties of these porins is instrumental to the development of new antibiotics. A common macroscopic feature of many OM porins is their ability to transition between functionally distinct open and closed states that regulate transport properties and rate. To obtain a molecular basis for these processes, we performed tens of microseconds of molecular dynamics simulations of *E. coli* OM porin, OmpF. We observed that large-scale motion of the internal loop, L3, leads to widening and narrowing of the pore, suggesting its potential role in gating. Furthermore, Markov state analysis revealed multiple energetically stable conformations of L3 corresponding to open and closed states of the porin. Dynamics between these functional states occurs on the time scale of tens of microseconds and are mediated by the movement of highly conserved acidic residues of L3 to form H-bonds with opposing sides of the barrel wall of the pore. To validate our mechanism, we mutated key residues involved in the gating process that alter the H-bond pattern in the open/closed states and performed additional simulations. These mutations shifted the dynamic equilibrium of the pore towards open or closed states. Complementarily, the mutations favoring the open/closed states lead to increased/decreased accumulation of multiple antibiotics in our whole-cell accumulation assays. Notably, porins containing one of the mutations favoring the closed state has previously been found in antibiotic resistant bacterial strains. Overall, our 180 *µ*s of simulation data (wild type and mutants) with concerted experiments suggests that regulation of the dynamic equilibrium between open and closed states of OM porins could be a mechanism by which Gram-negative bacteria acquire antibiotic resistance.

## Introduction

Antibiotic resistance is a major concern in treatment of bacterial infections. ^1–3^ Design of antibiotics targeting Gram-negative bacteria is particularly challenging due to the presence of an outer membrane (OM) containing lipopolysaccharide glycolipids. The OM provides a major permeation barrier against the uptake of various substrates including antibiotics.^2,4–11^ Due to the impermeability of the OM, essential nutrients for the bacteria typically diffuse into the cell through a variety of general-diffusion, *β*-barrel OM porins.^6^ Several OM porins have been shown to be also the main pathways for penetration of antibiotics into Gram-negative bacteria. ^6,12^

The general-diffusion OM porins are typically organized into trimeric *β*-barrel structures with multiple loops connecting individual *β* strands in each monomer. Of particular functional interest is a long internal loop (L3) that folds into the lumen of the monomeric *β*-barrels and forms a constriction region (CR) in the pore. The CR significantly narrows the pore and acts as a major barrier to substrate permeation.^13^ Additionally, the CR contains many charged and polar residues including acidic residues on the L3 loop, and a cluster of tyrosine and basic residues on two opposite sides of the barrel wall (termed here as Y- and B-face, respectively) (Fig. 1). The nature and organization of the B-face and Y-face residues will determine their interaction with the L3 loop, and therefore are expected to influence the dynamics of L3 and thus the permeation properties of the porin.

**Figure 1:**
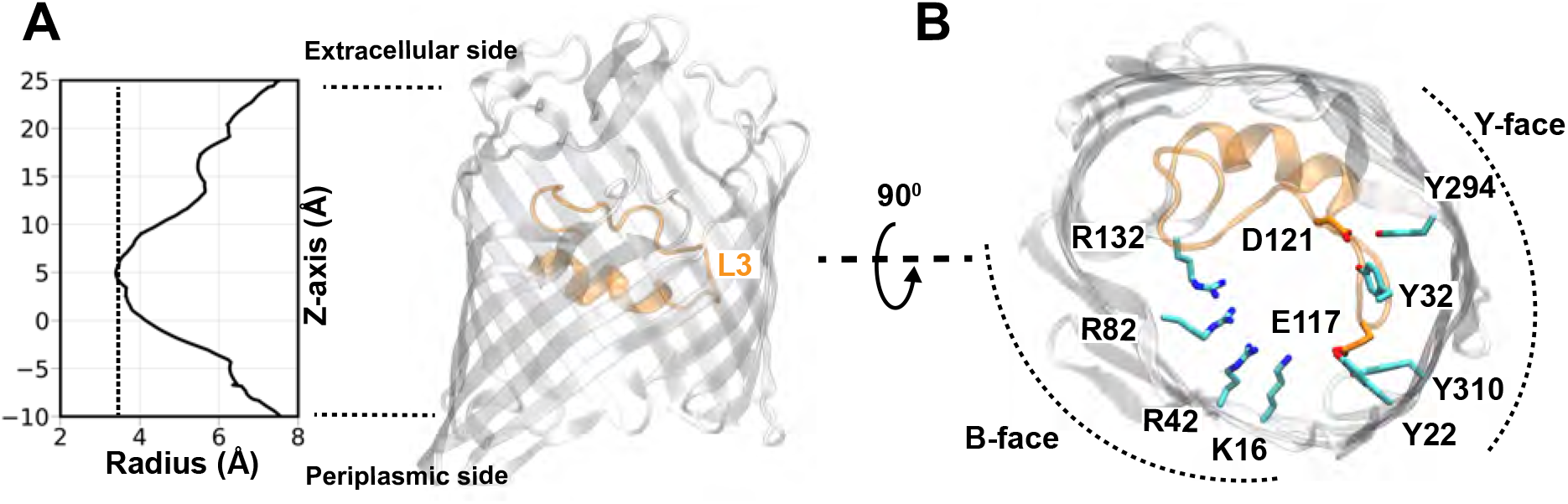
Structural features of OmpF in *E. coli*. (A) The monomeric radius profile (calculated using the program HOLE^28^) of the crystal structure of OmpF (PDB ID: 3POX^26^). A molecular representation of a single monomer of OmpF is shown highlighting the internal loop L3 (orange) that constricts the pore. (B) A top-down view of OmpF showing two acidic residues of L3 (D121 and E117) form hydrogen bonds with residues in the Y-face. A cluster of basic residues (B-face) on the opposite side of the Y-face is hypothesized to facilitate the movement of L3 to further narrow the pore.

A remarkable feature of many OM porins is their ability to undergo spontaneous conformational transitions between macroscopically distinct “open” and “closed” states. Gating motions in other membrane transport proteins are often active processes, coupled to and driven by an external energy input, such as the transmembrane voltage change.^14–18^ However, given the absence of energy in the OM, the energy provided by thermal fluctuations is likely the source to support spontaneous changes during the transition between conducting and non-conducting states in OM porins. ^19–24^ The most abundant OM porin in *Escherichia coli*, OmpF, is an ideal model to study such conformational changes; OmpF is known for spontaneously fluctuating between highly stable, conducting (open) and less stable, non-conducting (closed) states, as observed in electrophysiological measurements. ^21,22,25^ However, the molecular mechanism by which OmpF undergoes these spontaneous gating processes is still not well understood.

We expect that the internal loop, L3, might play an important role in gating the OmpF due to its location within the CR. Several of the acidic residues in L3, including E117 and D121, interact with tyrosine residues of the Y-face in the crystal structure of OmpF (PDB ID: 3POX) (Fig. 1(A)).^26^ It has been observed in a previous molecular dynamics (MD) study that L3 leaves the Y-face to transiently interact with the B-face thereby narrowing the pore. ^27^ These results led to the hypothesis that L3 movement could be responsible for gating of the pore. However, due to the short duration of the simulation and the nonphysiological conditions (vacuum) used in this computational study, structural support for this hypothesis is lacking.

As an indirect support for this hypothesis, mutation of B-face residues to uncharged or negatively charged residues showed an increase in substrate permeation,^29–31^ possibly as a result of reduced attraction of L3 to the B-face, shifting the equilibrium towards the open state. Furthermore, the crystal structure of a clinically relevant mutant of OmpF, G119D,^32^ suggested that adding negative charge to L3 can potentiate its attraction towards the B-face, thereby further reducing the pore size (Fig. S1) as compared to the wild-type, shifting therefore towards the closed state. This shift resulted in decreased permeability of the mutant to substrates such as carbohydrates and antibiotics.^32^ Strikingly, this mutant porin was shown to confer colicin N resistance to clinical strains of *E. coli*.^32^ The purpose of this study is to provide a comprehensive atomistic insight into the mechanisms controlling the dynamic equilibrium between open and closed states of OmpF using extensive MD simulations which are used to construct a Markov state model (MSM) for the process. MSMs are a class of model used to describe the long timescale dynamics of molecular systems and to obtain the thermodynamic and kinetic information about dynamic processes from the MD simulation data. ^33^ Using MSMs, we find that large-scale motion of L3 controls the dynamic equilibrium between conducting and non-conducting states of OmpF. Further analysis using the transition path theory^34^ revealed that transitions between open and closed states occur in two steps: 1) movement of E117 to the B-face to initiate the transition from the open state, 2) movement of D121 to the B-face that drives a large scale movement of L3 to mediate complete closure of the pore. Furthermore, simulations of charge-reversal mutants of B-face residues key in our proposed mechanism show a significant decrease in the probability of the closed state. This agrees with the increased accumulation observed for several antibiotics in our whole-cell assays in which we expressed the mutant porins. Furthermore, previous experiments also report increase in substrate permeability of these mutants.^29–31^ According to our model, these mutations reduce the attraction of E117 or D121 to the B-face thereby reducing the closed state probability, leading to an increased likelihood of the open state. Overall, our results provide a mechanistic understanding of how thermal motion of an internal loop control a dynamic equilibrium between open and closed states, therefore regulating permeability of OM porins.

## Results and Discussion

### Conformational dynamics of L3 mediate OmpF gating

To extensively sample multiple conformations of L3 that could lead to functionally distinct states, we performed ten independent replicas of MD simulation, each for 1 *µ*s, starting from the crystal structure of OmpF trimer (PDB ID: 3POX)^26^ embedded in a lipid bilayer. In multiple replica, the root mean squared deviation (RMSD) of L3 (backbone atoms) was observed to reach values greater than 4Å from the crystallized state, indicating a large-scale motion of this loop (Fig. 2A). This large-scale motion of L3 was accompanied with widening and narrowing of the CR in the pore, (Fig. 2B), suggesting its potential role in gating of OmpF. The structural deviation of L3 from its initial conformation, which was even observed in the early phase of a few replicas, was seemingly unexpected, because the crystal structure represents a thermodynamically stable configuration. However, since we performed our MD simulations at a more relevant biological temperature (310 K) ^35^ as opposed to the relatively low temperature (296 K) at which OmpF was crystallized at,^26^ the observed rapid structural deviation of L3 was reasonable.

**Figure 2:**
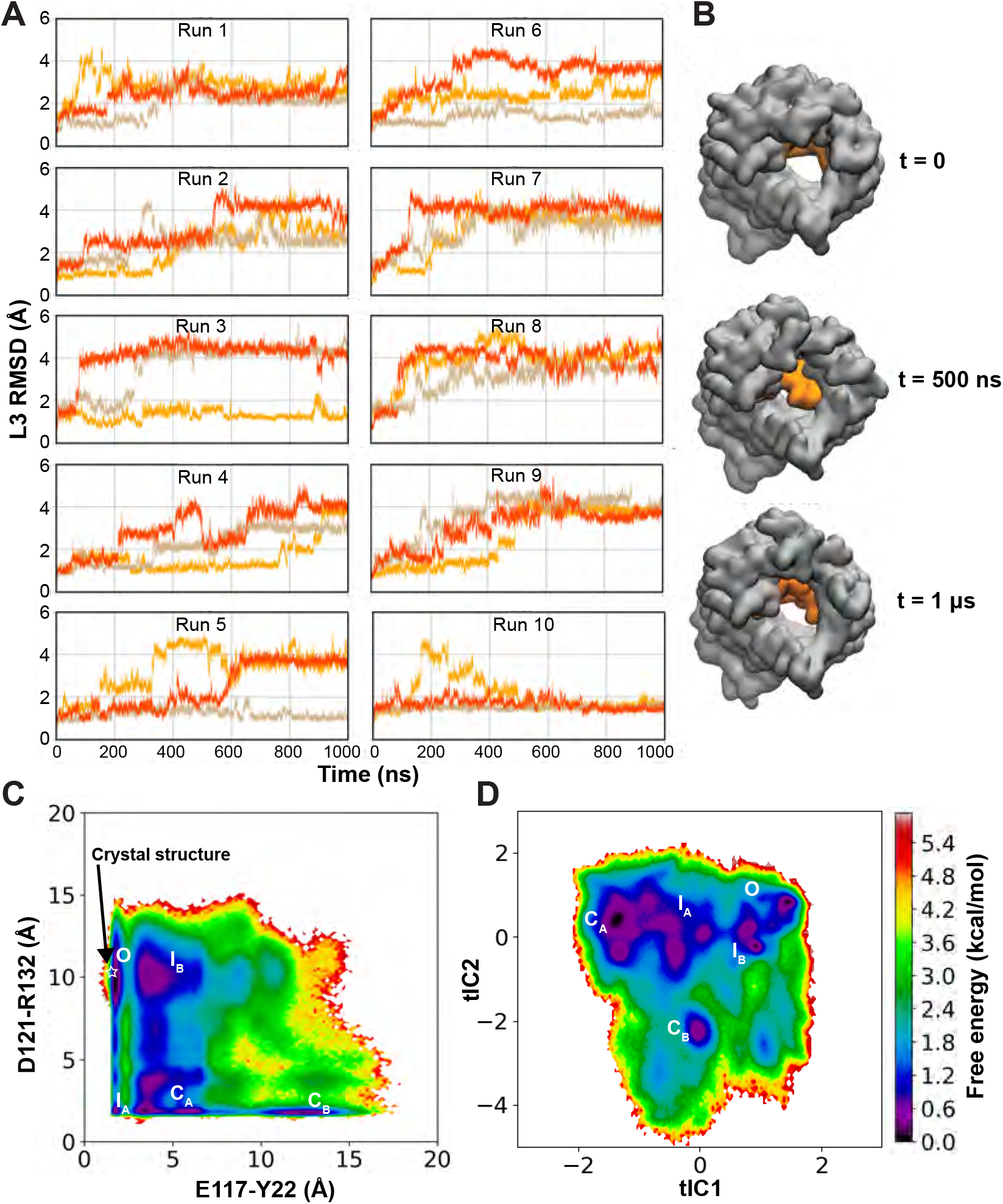
Dynamics and conformational landscape of L3. (A) Time evolution of root mean-squared deviation (RMSD) of loop L3 (after aligning the barrel residues to the crystal structure) in each monomer (shown in a different color) in all 10 replicas (Runs 1–10). (B) Conformation of L3 (shown in orange) of a single monomer at different time points along a representative trajectory of membrane-embedded OmpF. L3 appears to close and reopen the pore during the trajectory. (C) The free energy landscape for L3 dynamics in OmpF, reweighted by the stationary distribution, is projected onto the space formed by the E117-Y22 and D121-R132 closest distances (optimal indicators of the slowest process, more details in the Methods section). Conformational states corresponding to energetic minima are represented as *O, I*_*A*_, *I*_*B*_, *C*_*A*_, and *C*_*B*_, representing open, intermediate, and closed states (see text for details). The white star indicates the crystal structure of OmpF (PDB ID: 3POX). (D) Free energy landscape projected onto the top two tICA eigenvectors. The five conformational states (identified from the free energy projection onto E117-Y22 and D121-R132 distances) are mapped onto this surface.

To probe the key amino acids contributing to the loop’s dynamics and possibly gating of the pore, we calculated the root mean squared fluctuation (RMSF) of the C*α* atom of each residue in L3 (Fig. S2). We found that residues 116 to 123 are the most flexible with C*α* RMSFs greater than 2Å in at least one of the monomers of any simulation replica. The largest fluctuation was observed in the residue G120, located at the tip of the L3 loop, with a maximum RMSF of 5.2Å (Fig. S2). A high flexibility for G120 has also been reported in a previous MD simulation study.^36^

During our simulations, the dynamics of L3 residues resulted in differential hydrogen-bonding patterns with two opposite faces (Y and B-face) of the *β*-barrel wall. A total of 26 hydrogen bonds can be defined based on either (i) high occurrence probability (occurrence > 25%), or (ii) long lifetime (maximum lifetime > 50 ns), suggesting their importance in the transition of L3 between different conformational states.

To identify key conformational states of L3 and kinetics of state transitions, we constructed an MSM by featurizing our trajectory data using the distances of these 26 hydrogen bond pairs. Since these distances are uncorrelated between monomers (Fig. S3), indicating no significant monomer-monomer cooperativity, the trajectory of each monomer can be considered independently, resulting in an aggregate of 30 *µ*s (10 independent runs × 3 monomers × 1 *µ*s). We then performed time-lagged independent components analysis (tICA) on this data set for dimensionality reduction and kept the top 5 tICs, which sufficiently describe slow transitions of the system (Figs. S5 and S6). The tICA space was then discretized into 1,000 microstates, which were used to build an MSM with a lag time of 2 ns (Figs. S6 and S7, see Methods for more details).

To reveal key conformational states, we evaluated the free energy landscape, reweighted by the stationary distribution obtained from MSM. To identify physically meaningful metrics to project this landscape, we evaluated the correlation of each distance feature with the second eigenvector of the TPM (Fig. S8) since it has been established that features correlated to this eigenvector optimally describe the slowest transitions of the system. ^37^ We then projected the reweighted free energy landscape onto the features with the greatest positive or negative correlation (E117-Y22 and D121-R132 distances) (Fig. S8), corresponding to breaking and formation of hydrogen bonds, respectively. From the free energy landscape (Fig. 2C), we identified 5 distinct conformational states corresponding to energy minima. These 5 states also correspond to minima in the MSM reweighted free energy landscape projected onto the first two tICA eigenvectors (Fig. 2D), indicating that these states are not simply a result of the choice of representative metrics.

To evaluate the functional characteristics of these conformational states, we computed the pore radius profile of each state using the HOLE program^28^ (Fig. 3). Based on the calculated pore bottleneck radius, we identify distinct functional states: an open state (*O*) with the widest bottleneck radius of 3.0 ± 0.2Å and two closed states (*C*_*A*_, *C*_*B*_) with the narrowest bottleneck radii of 1.3 ± 0.5 and 1.4 ± 0.4 Å, respectively. Additionally, two intermediate states (*I*_*A*_, *I*_*B*_) with bottleneck radii of 2.4 ± 0.3 and 1.9 ± 0.6Å were also identified. Thus, we have shown that L3 dynamics indeed drive gating processes of the pore between open and closed state conformations.

**Figure 3:**
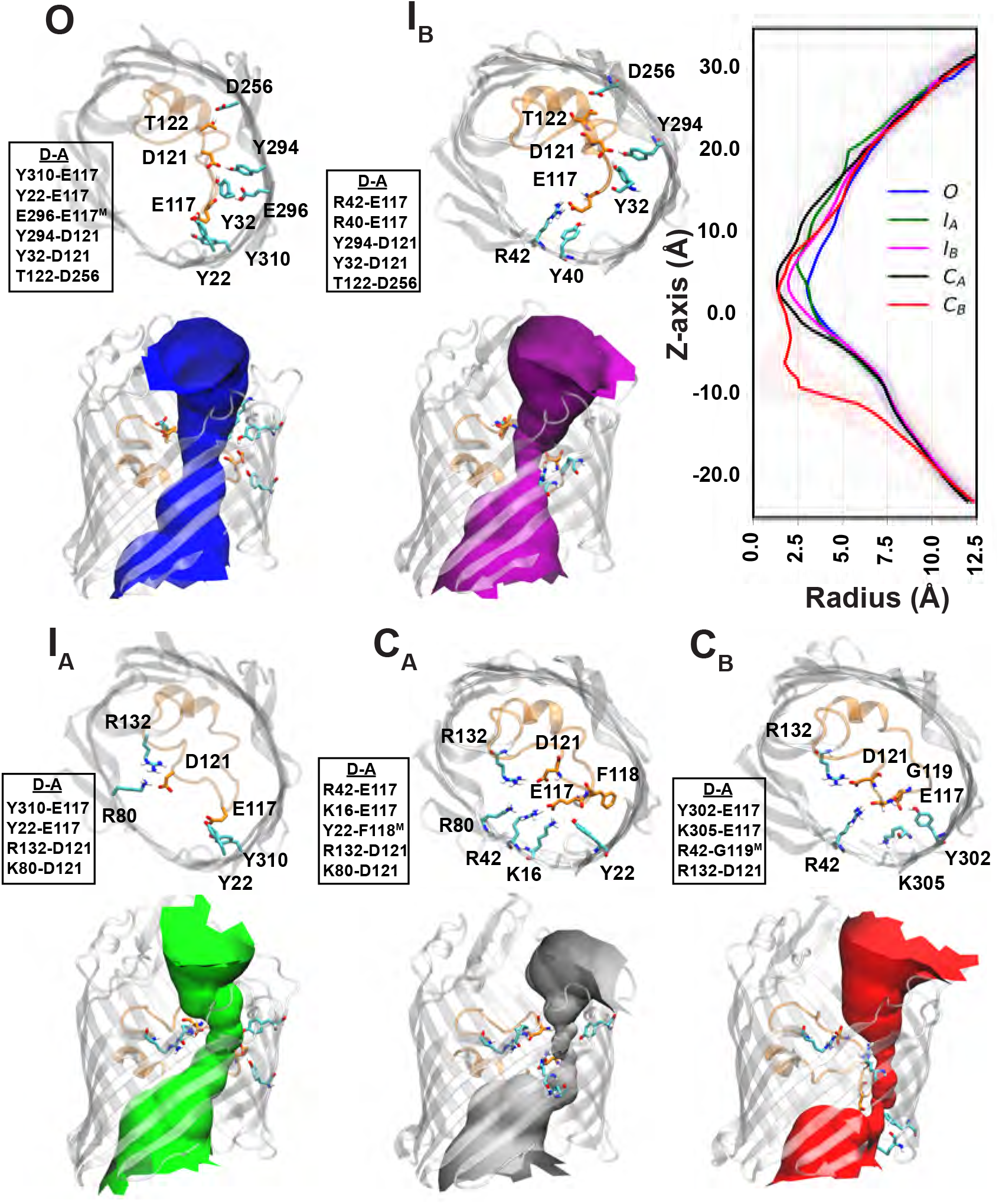
Structural characteristics of the conformational states in OmpF. The figure shows top-down snapshots of the five conformational states (*O, I*_*A*_, *I*_*B*_, *C*_*A*_, and *C*_*B*_) highlighting hydrogen bonds with > 20% occurrence probability between the most fluctuating residues of L3 (Fig. S2) and the barrel residues. The abbreviation “D-A” here refers to the Donor-Acceptor pair of the hydrogen bond. The ensemble distribution of distances of each hydrogen bond for the conformational states can be seen in Fig. S9. The pore profiles (determined using the program HOLE^28^) of the conformational states are depicted with different colors in the side view of the pore. The mean (solid line) and standard deviation (shaded region) of the radius profile calculated using HOLE^28^) are shown for each conformational state.

### Distinct gating mechanisms mediated by acidic residues, E117 and D121

Two distinct gating processes play major roles in the observed dynamics of L3 in OmpF: *O* to *C*_*A*_ transition, with a mean first passage time (MFPT) of 4.6 ± 0.1 *µs*, and the significantly slower *O* to *C*_*B*_ transition with an MFPT of 40.5 ± 2.0 *µs*. To obtain mechanistic insight on the two gating processes, pathways of conformational transitions from *O* to *C*_*A*_ and from *O* to *C*_*B*_ were identified separately using transition path theory (TPT).^34,38,39^ The two gating processes were analyzed by grouping their pathways according to conformational states visited, which allowed us to identify key intermediates in gating. We found that in half of the pathways involved in both gating processes, initial transitions are from *O* to the intermediate state *I*_*B*_ (Figs. 3 and 4). These transitions are primarily mediated by movement of E117 a short distance (∼ 2.5 Å) from the Y-face to the B-face where it interacts with R42 and Y40. In comparison, direct *O* to *C*_*A*_ transitions are mediated by movement of both E117 and D121 from the Y-face to the B-face. These transitions have a lower likelihood, since D121 needs to travel a larger distance (∼ 10 Å) to reach its new interaction partners, R132 and R80 (Figs. 2 to 4). Pathways involving an initial movement of only D121 (and not E117) to the B-face (*O* to *I*_*A*_ transition) have negligible total fluxes.

**Figure 4:**
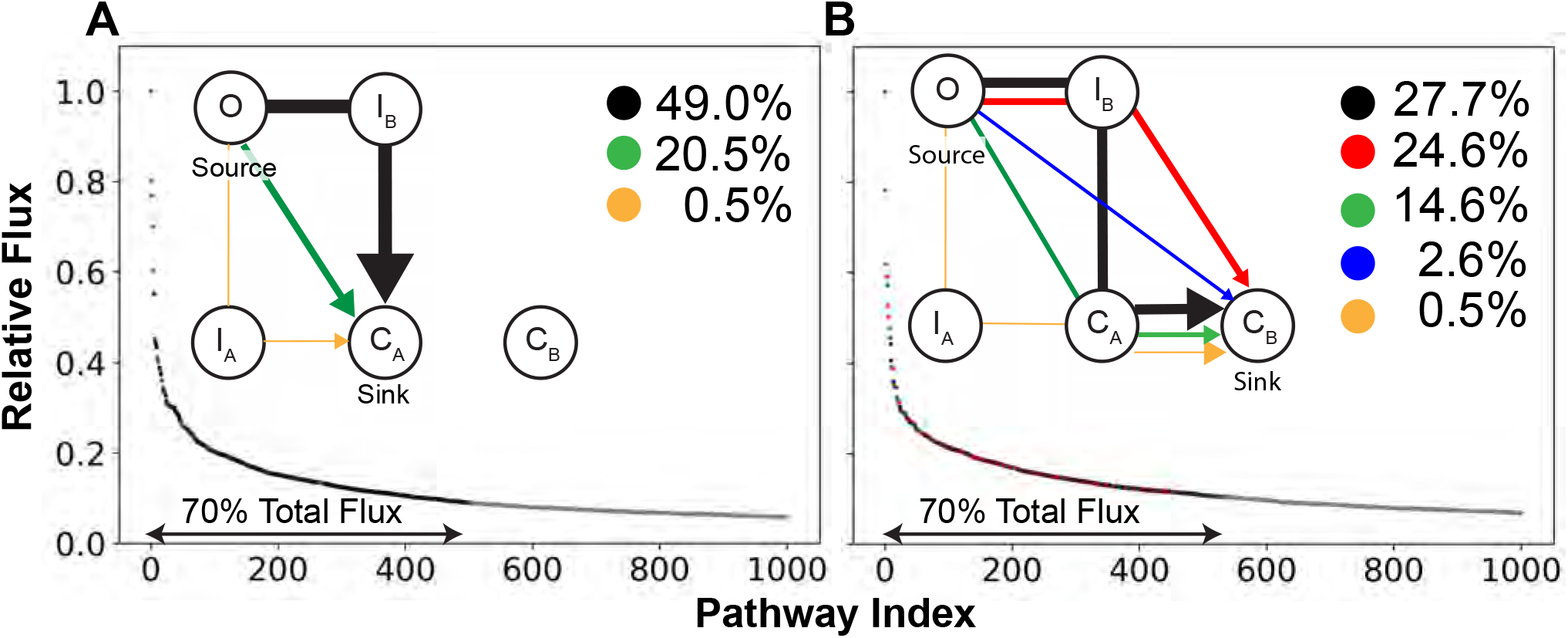
Dominant gating transition pathways in WT-OmpF. The pathways from *O* to *C*_*A*_ (A) and from *O* to *C*_*B*_ (B) identified from transition path theory are ranked according to their flux relative to the highest flux path. The paths representing top 70% of the total flux are grouped according to the intermediate conformational states visited, and color coded black, red, green, blue, or orange according to the pathway group. The flux sum for all paths belonging to each pathway group is depicted in the legend. (*Inset*) Schematic of each pathway group for *O* to *C*_*A*_ (A) and *O* to *C*_*B*_ (B) transitions. The width of the arrow approximates the flux sum of each group.

After reaching the intermediate state *I*_*B*_ (observed in half of the transitions), movement of D121 from the Y-to B-face triggers a large scale motion of the entire L3 loop towards the B-face and completes closure of the pore (transitions to either *C*_*A*_ or *C*_*B*_) (Fig. 3). This D121-mediated loop movement is preserved in transitions from *I*_*B*_ to either *C*_*A*_ or *C*_*B*_. However, transition to *C*_*B*_ requires additional movement of E117 (along the pore axis and towards the periplasmic space) to interact with K305 and Y302 located below the B-face (Fig. S4A), which elongates the narrow region of the pore. The downward movement of E117 is accompanied by partial unfolding of the *α*-helical portion of L3, and conformational restriction of the backbone torsion of a proline residue, P116 at the tip of the loop (Fig. S4). This offers a possible explanation for the slow timescale of the gating processes involving the formation of *C*_*B*_. Overall, our analysis suggests that pore closure generally occurs in two key steps: (1) E117 movement to the B-face (R42 and K16) mediating the initial transition out of the *O* state, and (2) D121 movement to the B-face (R132) mediating large-scale motion of L3 leading to closure of the pore.

Our mechanism for gating processes could be generalized to OM porins of *Enterobacteriaceae*, such as OmpC and PhoE in *E. coli*, OmpK36 of *K. pneumoniae*, and OmpE36 from *E. cloacae*, since these porins share a similar architecture with OmpF and conserve key residues involved in gating (Fig. S10). However, our mechanism should not be generalized to porins with multiple internal loops such as the OccK and OccD family in *P. aeruginosa* since their gating might involve the motion of multiple loops.^40,41^

### Mutations regulating equilibrium dynamics of the loop determine OM permeability to antibiotics

Many of the key residues in the identified gating mechanism are shown to be important for permeation through OM porins of *Enterobacteriaceae* in our own and previous experiments. In our whole-cell assays (details below), we compared the accumulation of various antibiotics (nalidixic acid, tetracycline, and enoxacin) in *E. coli* BW26678 (parental strain) with a strain containing a point mutation of a key B-face residue (R132) to a negatively charged side chain in OmpF, *E. coli* OmpF^R132D^. We found a statistically significant increase in the accumulation of the two negatively charged antibiotics, nalidixic acid and tetracycline, for cells expressing the mutant (Fig. 6). However, accumulation values of the zwitterionic enoxacin (at physiological pH) were not significantly altered, possibly because of the accumulation of enoxacin through WT-ompf is already saturated (Note substantially higher accumulation values for this compound in Fig. 6). The higher accumulation of other zwitterionic antibiotics has also been described before^42^ and might be due to differential engagement of these drugs with other uptake/efflux mechanisms.

**Figure 5:**
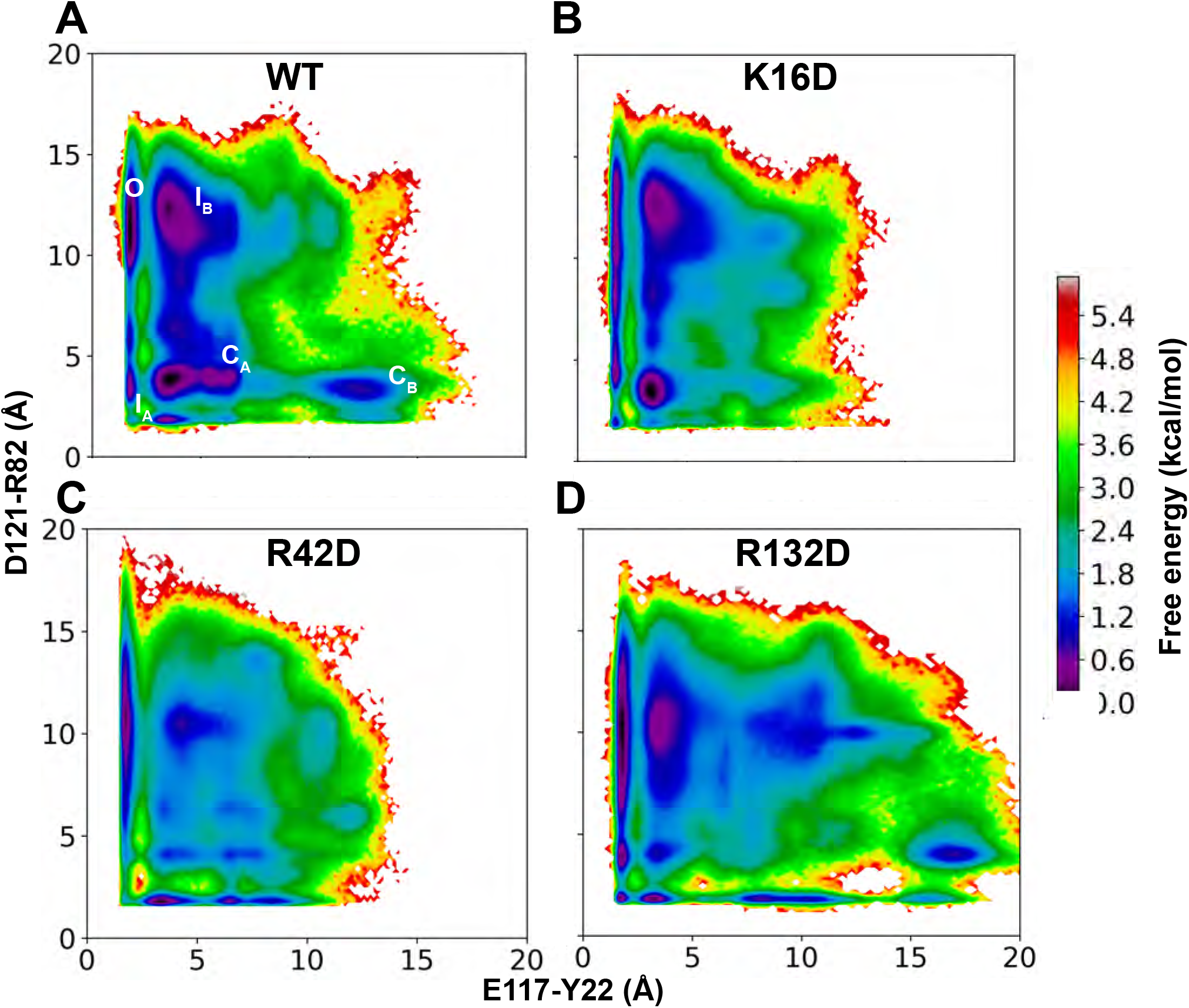
Conformational landscape of L3 in the wild-type (WT), K16D, R42D and R132D-OmpF simulations. (A) The free energy landscape for the WT-OmpF, reweighted by the stationary distribution, is projected onto the E117-Y22 and D121-R82 distance features. The conformational states identified in Fig. 2C,D are mapped onto this landscape. (B, C, D) Free energy landscapes for K16D, R42D and R132D-OmpF. The free energy error for each system can be seen in Fig. S11.

**Figure 6:**
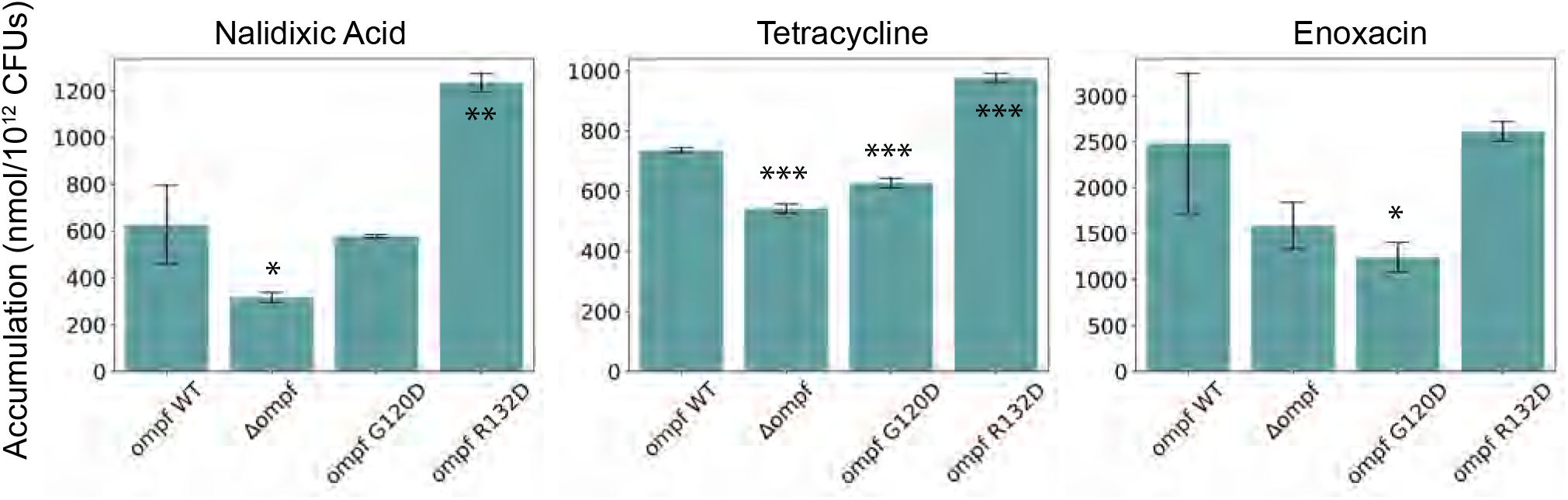
Accumulation values of three different antibiotics in WT *E. coli* BW26678 (parental strain) and three modified strains, including complete deletion of the *ompF* locus, *E. coli* OmpF^G120D^, and *E. coli* OmpF^R132D^. Accumulation values are reported in nmol per 10^12^ colony-forming units (CFUs). Data shown represent the average of three independent experiments. Error bars represent the standard deviation of the data. Statistical significance was determined by a two-sample Welch’s t-test (one-tailed test, assuming unequal variance) relative to accumulation values in the WT strain. Statistical significance is indicated with asterisks (* p < 0.05, ** p < 0.01, *** p < 0.001).

Previous experiments including liposome swelling assays, radiolabeled substrate uptake assays and antibiotic susceptibility measurements have also shown that mutation of key B-face residues in OmpF to uncharged or negative residues (R42S, K16D, R132A, and R132D) significantly increase permeation of various substrates including carbohydrates and antibiotics.^29–31^ Additionally, mutation of a conserved basic residue, K16, in OmpC to a negatively charged residue (K16D) in electrophysiology measurements showed significant increase in the probability of open, conducting states.^20^ The rationale previously given for the increased substrate permeation was that these mutations decreased the bulkiness of the CR thus increasing the pore radius, or altered substrate-protein interactions to facilitate permeation.^29,31^ Our study, however, suggests an alternative hypothesis: these mutations increase permeation by altering the OmpF gating dynamics. In particular, the K16D or R42S mutations would hinder the initial transition of OmpF out of the open state by reducing E117 attraction to the B-face. The R132A or R132D mutations would hinder the large-scale motion of L3 by reducing D121 attraction to the B-face. Both set of mutations would decrease the probability of pore closure transitions, leading to an increased likelihood of the open state.

To test this hypothesis, we built three mutant systems, R42D, K16D and R132D. in all three monomers of the crystal structure of OmpF trimer (PDB ID: 3POX).^26^ Using these three mutant structures as starting points, ten independent 1 *µ*s simulations were performed for each mutant. Simulation trajectories were then used to build an independent MSM for each mutant system following the same procedure as for the WT. We evaluated the free energy landscapes, reweighted by the stationary distribution obtained from MSM, for each mutant. To compare the results, free energy landscapes of WT and mutant systems were projected onto the distance features, E117-Y22 and D121-R82 (Fig. 5). D121-R82 was used as a distance feature instead of D121-R132, since the R132D mutation abolishes a favorable interaction between these sites. In the K16D and R42D systems, an increased energetic barrier compared to the WT system was observed for the *O* to *I*_*B*_ transition (Fig. 5B,C). This supports the hypothesis that a negative charge at residue 16 or 42, the interaction sites of E117 upon transition to the B-face in WT, hinders the initial transition out of the *O* state (Figs. 2 and 4). In the R132D system, the energetic barrier for the *O* to *I*_*B*_ transition was not substantially altered (Fig. 5D) since this mutation does not affect the movement of E117. Regardless, in each mutant system, at least one closed state probability was significantly lowered compared to the WT (Fig. 5), in accordance with our hypothesis that both D121 and E117 movements are necessary for closed state formation.

In general, the proposed model suggests that the permeability of OmpF depends on the dynamic equilibrium between the open and closed conformations of the pore. This is in contrast to the typical understanding of the pore as a rigid body permeating substrates.^43,44^ An example of how a shift in dynamic equilibrium impacts permeability has been previously captured in the clinically relevant, G119D mutant of OmpF. The crystal structure of the G119D-OmpF^32^ shows that the introduced negative charge on L3 increases attraction of the loop to the B-face, thus stabilizing the pore in a narrower state than in the WT structure (Fig. S1). The loop movement reduces the pore radius compared to the open WT crystal structure (Fig. S1). Further relaxing the mutant crystal structure in microsecond-scale MD simulations (in multiple replicates) followed by MSM analysis, we find additional shifts in equilibrium towards the closed state (Fig. S14). We identify two structurally distinct closed states on the conformational landscape of L3 (bottleneck radii of 0.9 ± 0.3 and 1.8 ± 0.3 Å) for this mutant. These shifts in equilibrium relate to the significantly decreased permeability of the G119D mutant to substrates such as carbohydrates and antibiotics.^32^ Strikingly, the reduced permeability of this mutant porin was shown to confer colicin N resistance to clinical strains of *E. coli*.^32^

To generalize the idea that a negative charge can shift L3 dynamics, we built another mutant system, G120D, and applied our approach of microsecond-scale MD simulations (in multiple replicates) and MSM analysis to this mutant. Each energetically stable state found for G120D exhibits a narrower pore (bottlenecks of 2.3 ± 0.6, 1.2 ± 0.6 and 1.0 ± 0.6 Å, respectively, for the three identified stable states) than the open state of WT-OmpF (Fig. S15). To verify experimentally the potential effect of the G120D mutation on antibiotic permeation, as suggested by our simulations, we compared the accumulation of the three antibiotics in *E. coli* BW26678 (parental strain) with a strain containing the G120D point mutation in OmpF, *E. coli* OmpF^G120D^. We found a statistically significant decrease in the accumulation of tetracycline and enoxacin for cells expressing the mutant (Fig. 6). However, we did not observe a significant decrease in accumulation of nalidixic acid, possibly because nalidixic acid is markedly smaller than the other two antibiotics and could still permeate through narrower porins. In general, our study suggests that regulation of the dynamic equilibrium between open and closed states of OM porins could be a mechanism by which Gram-negative bacteria become resistant to antibiotics.

## Concluding Remarks

Outer membrane (OM) porins form the main pathway for exchange of material between the environment and the intracellular compartment in Gram-negative bacteria. Due to their central role in transport, OM porins are a key factor in the mechanism of antimicrobial resistance of a large number of bacteria. Using extensive MD simulations to construct an MSM, we characterize the gating mechanisms in OmpF, the archetypal OM porin in *E. coli*. From the MSM, combined with structural analysis of the involved states, we find that large-scale motion of a folded loop, L3, controls the transition of OmpF between open and closed states. The gating mechanisms captured during multi-microsecond simulations of membrane-embedded OmpF is primarily mediated by: (1) shift of E117 to a cluster of basic residues (B-face) causing an initial departure from the open state, and (2) movement of D121 to the B-face driving large-scale motion of L3 and completing closure of the pore. The observed mechanism may be generalized to several other Gram-negative porins with similar architecture and conserved key residues involved in gating (Fig. S10). Supported by the increased antibiotic accumulation inside *E. coli* in our experiments and increased permeability to carbohydrates and antibiotics reported in previous experiments,^29–31^ *µ*s-scale simulations of charge-reversal mutants of key B-face residues show a decreased probability for the closed state. From the results, we provide a model indicating that the permeability of OM porins depends on the dynamic equilibrium between the open and closed conformations of the pore. This model could offer a new perspective on the mechanisms by which OM porins mediate antibiotic resistance in Gram-negative bacteria.

## Methods

### Preparation and simulation of membrane-embedded OmpF

We used the X-ray structure of OmpF trimer (PDB ID: 3POX),^26^ including all the crystal water molecules, as a starting point for our simulations. In each monomer, residues E296, D312, and D127 were protonated in accordance with previous studies.^45–47^ The OmpF trimer was embedded in a symmetric membrane composed of 1,2-dimyristoyl-sn-glycero-3-phosphocholine (DMPC) lipid molecules. Usage of an OM composition containing lipopolysaccharides (LPS) is not considered, since the membrane composition is unlikely to influence the dynamics of L3 as this internal loop is fully internalized in the protein fold and not exposed to the membrane. The protein-DMPC system was then solvated with TIP3P water^48^ and buffered in 0.15 M NaCl. Each step of the membrane building process was carried out using the Membrane Builder module of CHARMM-GUI.^49^ The final system contained ∼140,000 atoms with dimensions of 120 × 120 × 100 Å^3^. Then, 10 independent molecular dynamics simulations were run. In each simulation, the prepared system was minimized using the steepest descent algorithm for 2,000 steps, followed by an initial equilibration of 5 ns, during which the protein heavy atoms were harmonically restrained using a force constant of 5 kcal mol^−1^ Å^−2^. Then, 1 *µ*s of unrestrained production simulation was performed for each replica.

### Molecular dynamics (MD) simulation protocol

MD simulations in this study were performed using NAMD^50,51^ utilizing CHARMM36m^52^ and CHARMM36^53^ force field parameters for proteins and lipids, respectively. A timestep of 2 fs was used in all simulations, and periodic boundary conditions were employed in all three dimensions. Bonded and short-range nonbonded interactions were calculated every timestep. The particle mesh Ewald (PME) method^54^ was used to calculate long-range electrostatic interactions every 4 fs with a grid density of 1Å^−3^. A force-based switching function was employed for pairwise nonbonded interactions starting at a distance of 10Å with a cutoff of 12 Å. Pairs of atoms whose interactions were evaluated were searched and updated every 20 fs. A cutoff (13.5 Å) slightly longer than the nonbonded cutoff was applied to search for the interacting atom pairs. Constant pressure was maintained at a target of 1 atm using the Nosé-Hoover Langevin piston method. ^55,56^ Langevin dynamics maintained a constant temperature of 310K with a damping coefficient, *γ*, of 0.5 ps^−1^ applied to all atoms. Simulation trajectories were collected every 10 ps.

### Markov state model construction

We used our trajectory dataset to construct a Markov state model (MSM) using pyEmma, ^57^ which enabled us to obtain kinetic and thermodynamic information about the system. To build the MSM, first the trajectory dataset was featurized using 26 residue-residue distance pairs with significant hydrogen bonding (occupancy greater than 25% during the simulations, or a maximum lifetime greater than 50 ns in at least one of the monomers of any replica) between the highly fluctuating residues of L3 (residues 116-123) and the barrel wall. A hydrogen bond was counted between an electronegative atom with a hydrogen atom (H) covalently bound to it (the donor, D), and another electronegative atom (the acceptor, A), if the D-A distance is less than 3Å and the D-H-A angle is greater than 120°. For these residue pairs, the minimum distance (in each frame of every trajectory) between any donated H and any A atom was used to create the MSM feature space. Since the distance pairs are uncorrelated between monomers (Fig. S3), we considered each monomer as an independent trajectory, giving us an aggregate trajectory data of 30 *µ*s (10 independent runs × 3 monomers × 1 *µ*s).

To remove redundant information within the feature space and identify the slowest reaction coordinates, time-structure based independent component analysis (tICA) was used to reduce the dimensionality of the feature space (*X*(*t*)) to the eigenvectors of an autocovariance matrix, ⟨*X*(*t*)*X*^*T*^ (*t*+*τ*)⟩, with a lag time, *τ*=1 ns.^37,58–60^ It is important to choose an optimal number of tICA eigenvectors since an MSM built using too many eigenvectors would have microstates with low statistical significance due to finite sampling error.^61^ We found that the first seven tICA eigenvectors are sufficient to construct the MSM because only the distribution of these eigenvectors significantly differed from the normal distribution (Fig. S5). Further statistical analysis using an MSM scoring method, VAMP-2 score,^62^ discussed further in the next section, showed that the quality of an MSM does not significantly improve when using more than five tICA eigenvectors (Fig. S6). Thus, we chose to reduce the number of eigenvectors to five in our study.

The conformational space was then discretized into multiple microstates using k-means clustering. To choose the number of microstates to use in the model, we used the VAMP-2 score, ^62^ to evaluate the quality of MSMs built with different numbers of microstates. The VAMP-2 score converged when using five tICA eigenvectors and 1,000 microstates (Fig. S6); thus, we used this parameter set to build our MSM.

Then, a transition probability matrix (TPM) was constructed by evaluating the probability of transitioning between each microstate within a lag time, *τ*. To choose an adequate lag time to construct a TPM that ensures Markovian behavior, multiple TPMs were first created using multiple maximum-likelihood MSMs with different lag times. The implied timescales 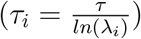 were evaluated for each of these transition matrices, and saturation was observed at *τ* = 2ns (Fig. S7). Thus, we built our final TPM using a maximum likelihood MSM with a lag time of 2 ns. This final TPM is symmetrized using a maximum likelihood approach to ensure detailed balance.^57^ This step did not significantly change the raw TPM (Fig. S12), indicating that the initial sampling was done under dynamic equilibrium conditions.

To identify physically meaningful metrics for projecting the free energy of the gating process, we used a protocol described by Pérez-Hernández *et al*. to choose the metric with greatest correlation to the second eigenvector of the TPM.^37^ The normalized correlation between the second eigenvector of the TPM and each of the 26 residue-residue distance pairs was evaluated as follows:

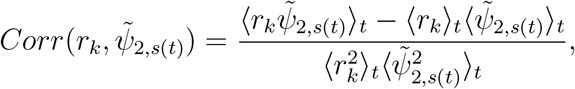

where *r*_*k*_ is the *k*_*th*_ residue-residue distance, 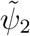 is the second eigenvector of the TPM, *s*(*t*) is the trajectory of microstates, and ⟨⟩_*t*_ is the time average. The E117-Y22 and D121-R132 distances were chosen for this purpose, as these features showed the greatest positive and negative correlations with the second eigenvector, respectively (Fig. S8). Using these features, we projected the free energy landscape weighted with the stationary distribution obtained from the MSM (Fig. 2C,D). The free energy landscape using the raw trajectory data, unweighted by the stationary distribution, is very similar to the weighted landscape (Figs. 2 and S13), indicating that the initial sampling used to build the MSM was sufficient to mitigate any sampling bias. To determine the error of the free energy landscape, we first used bootstrapping to alter the TPM, and to create new free energy landscapes. The error was then determined in every bin of the landscape. We used the free energy landscape to lump our microstates into 5 macrostates depending on whether the microstate physically lies within a free energy minima (defined using an energy cutoff of 1.2 kcal/mol) shown in Fig. 2C,D. Macrostates are classified according to their pore bottleneck radius (Fig. 3) leading to: an open state (*O*), two intermediate states (*I*_*A*_, and *I*_*B*_) and two closed states (*C*_*A*_, and *C*_*B*_).

### Transition path theory

To obtain kinetic information about gating processes, the mean first passage times (MFPTs) for the *O*-*C*_*A*_ and *O*-*C*_*B*_ transitions were evaluated. The uncertainty in the MFPT was evaluated using a Bayesian estimated MSM, implemented in pyEmma.^57^ The transition path theory module in pyEmma^57^ was used to identify conformational transitions involved in the gating process. This step was done by choosing the *O* and *C*_*A*_/*C*_*B*_ states as the source and sink, respectively, and identifying the pathways connecting them.

### Construction of *E. coli ompF* mutants

Strains carrying various *ompF* alleles were constructed in three steps. All strains, plasmids, primers and gBlocks used in these constructions are shown in Tables S1 to S4. Initially, a strain carrying a complete deletion of the *ompF* locus (Δ*ompF*8897::*cat*) was constructed by *λ*-red-mediated recombination as described elsewhere.^63^ To do this, BW26678 was transformed to Cm^*R*^ using a PCR product obtained using primers ompF-cloningF and ompF-cloningR with pKD3 as the template, creating WM8897. The Δ*ompF*8897::*cat* allele removes the entire *ompF* coding sequence and 323 base-pairs upstream of the start codon containing the promoter and regulatory sequences. Second, a series of plasmids carrying various *ompF* alleles were constructed using the pAH144^64^ as the vector. This plasmid encodes resistance to streptomycin and spectinomycin (Strep/Spec^*R*^) and can be inserted into the host chromosome in single copy at the Hong Kong phage attachment site (*att*-HK). Initially we constructed pMEM501, which carries the WT *ompF* gene and 610 upstream base-pairs, including the promoter and all known regulatory sequences. Plasmid pMEM501 was then modified by replacement of an appropriate internal restriction endonuclease fragment with a synthetic DNA fragment carrying the desired mutations. Finally, each of these plasmids was inserted into the chromosome of WM8901 by selection for Strep/Spec^*R*^. The inserts of all plasmids were verified by DNA sequencing. All strains were verified by PCR, including DNA sequencing of the PCR product, to confirm the presence of the Δ*ompF*8897::*cat* allele and the correct plasmid inserted in single copy.

### Accumulation assay protocol

The accumulation assay was performed in triplicate as outlined elsewhere.^12,65^ A 5mL overnight culture was diluted into 250mL of fresh lysogeny broth (LB) and grown at 37°C with shaking to an optical density (OD_600_) of 0.55-0.60. Once grown to mid-log phase, 200mL of culture was pelleted at 3,220 r.c.f. for 10 minutes (at 4°C). The supernatant was discarded and cells resuspended in 40 mL phosphate buffered saline (PBS), pelleted as before, and resuspended in 8.8 mL PBS. Cells were aliquoted into 1.7mL Eppendorf tubes each with 875 *µ*L and incubated with shaking at 37°C for 5 minutes to equilibrate cells. Colony forming units (CFUs) were determined by a calibration curve. These time points were short enough to minimize metabolic and growth changes (no changes in OD_600_ or CFU count observed). Cells were treated with 50 *µ*M compound (8.75 *µ*L of 5mM compound stock) for 10 minutes at 37°C with shaking. After incubation, 800 *µ*L of culture was layered over 700 *µ*L cold silicone oil (9:1 AR20/Sigma High Temperature, cooled to -78°C) and cells pelleted at 13,000 r.c.f. for 2 minutes at room temperature to separate supernatant and extracellular compound from bacterial cells. The supernatant and oil were removed by pipette and the cell pellet was resuspended in 200 *µ*L MilliQ water. Samples were subjected to three freeze-thaw cycles of alternating 3 minute incubation periods in liquid nitrogen (−78°C) and a 65°C water bath. Lysed cells were pelleted at 13,000 r.c.f. for 2 minutes and 180 *µ*L of supernatant were collected. Cell pellets were washed in 100 *µ*L methanol, vortexed, and pelleted at 13,000 r.c.f. for 2 minutes. After pelleting, 100 *µ*L of supernatant was collected and combined with previous supernatants. Remaining debris were removed through centrifugation at 20,000 r.c.f. for 10 minutes at room temperature. Supernatants were analyzed with the QTRAP 5500 LC/MS/MS system (Sciex) in the Metabolomics Laboratory of the Roy J. Carver Biotechnology Center, University of Illinois at Urbana-Champaign. Software Analyst 1.6.2 was used for data acquisition and analysis. The 1200 Series HPLC System (Agilent Technologies) includes a degasser, an autosampler and a binary pump. The liquid chromatography separation was performed on an Agilent Zorbax SB-Aq column (4.6mm × 50 mm; 5 *µ*m) with mobile phase A (0.1% formic acid in water) and mobile phase B (0.1% formic acid in acetonitrile). The flow rate was 0.3mL min^−1^. The linear gradient was as follows: 0–3 min: 100% A; 10–15 min: 2% A; 16–20.5 min: 100% A. The autosampler was set at 15°C. The injection volume was 1 *µ*L. Mass spectra were acquired under positive electrospray ionization with a voltage of 5,500 V. The source temperature was 450°C. The curtain gas, ion source gas 1 and ion source gas 2 were 33, 65 and 60 psi, respectively. Multiple reaction monitoring was used for quantitation with external calibration. All compounds evaluated in biological assays were ≥ 95% pure, assessed by NMR and LC-MS.

## Acknowledgements

This research is supported by National Institutes of Health grants R01-AI136773 (to P.J.H. and E.T.), R01-HL131673 (to E.T.), and P41-GM104601 (to E.T.). D.S. acknowledges support from National Science Foundation grant MCB-1845606. Simulations in this study have been performed using allocations at National Science Foundation Supercomputing Centers (XSEDE grant number MCA06N060) and the Blue Waters Petascale Computing Facility of National Center for Supercomputing Applications at University of Illinois at Urbana-Champaign, which is supported by the National Science Foundation (awards OCI-0725070 and ACI-1238993) and the state of Illinois.

## Author contributions statement

A.K.V., N.H., P.C.W., D.S., W.W.M., P.J.H., and E.T. designed research; A.K.V. and N.H. performed the simulations and analyzed data; R.J.U., M.E.M., W.W.M. performed the experiments and analyzed data; and A.K.V., N.H., D.S., and E.T. wrote the paper. All authors read and contributed to finalizing the paper.

## Data availability statement

The data that support the findings of this study are available from the corresponding author upon reasonable request.

