## Supplementary Information for "Investigation of gating in outer membrane porins provides new perspectives on antibiotic resistance mechanisms"

### Supporting Information

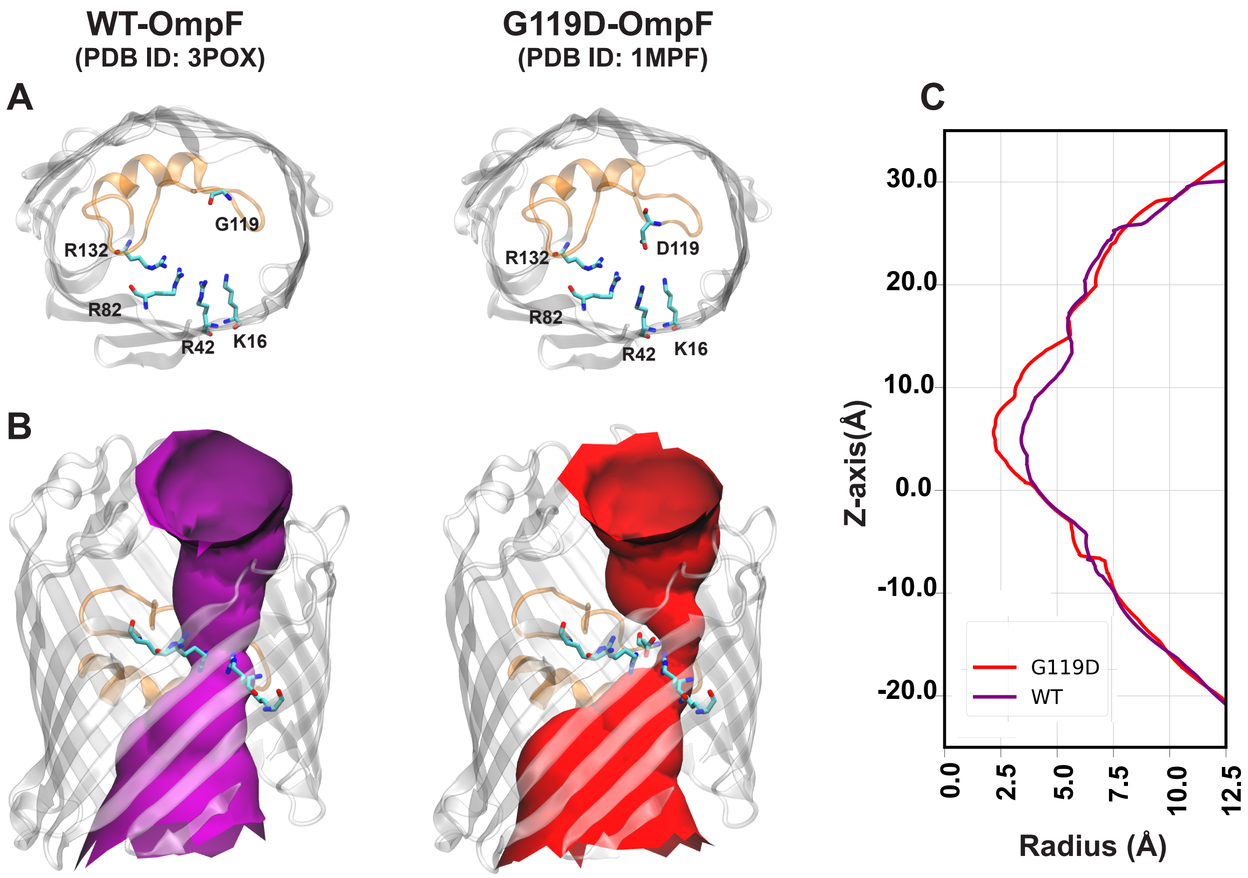


Figure S1: (A) Top-down view of WT-OmpF and G119D-OmpF crystal structures, highlighting position 119 and the B-face residues. (B) Pore profile (determined using the program HOLE[28]) in WT-OmpF and mutant G119D-OmpF crystal structures. (C) Radius profile of the pore calculated using HOLE[28] for each structure.

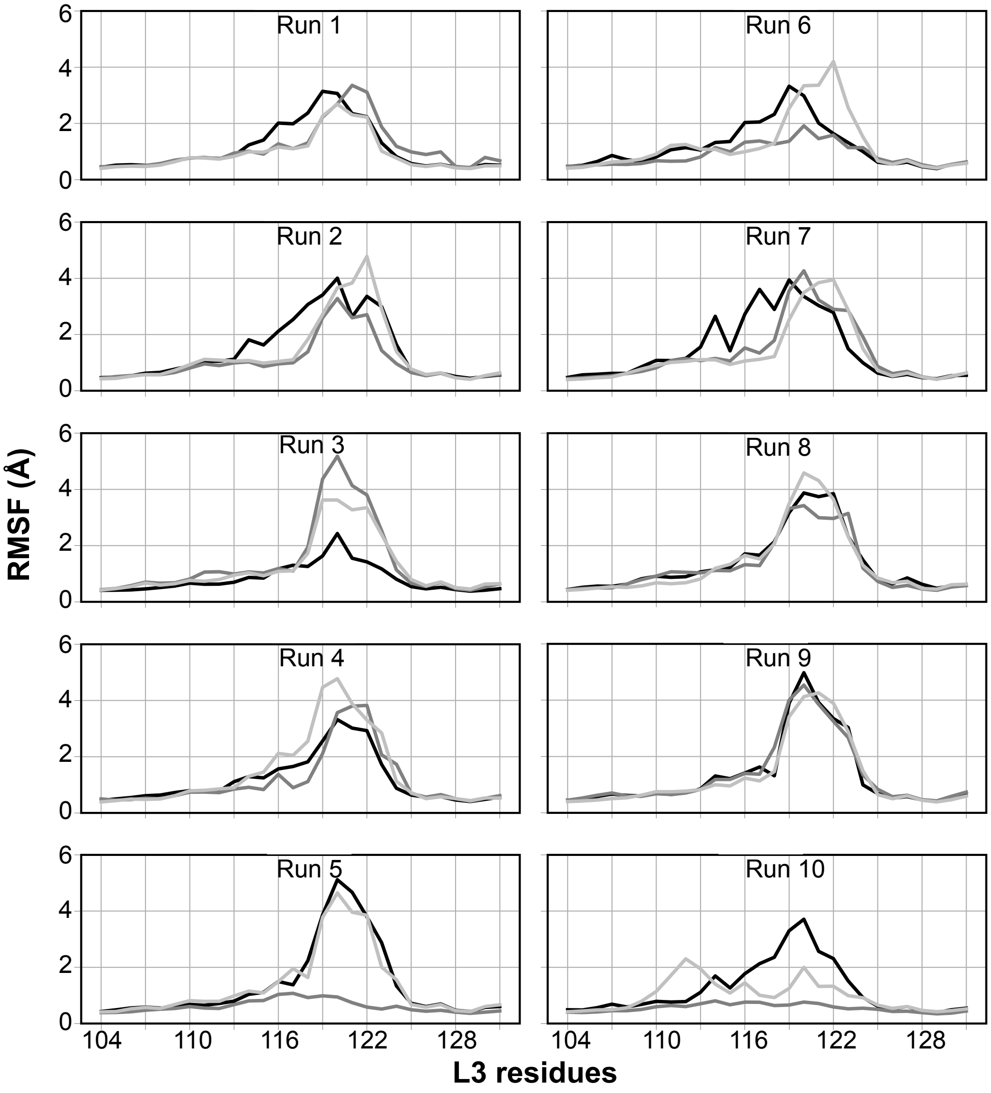


Figure S2: Root mean-squared fluctuations of L3 residues in each monomer in all 10 replicas.

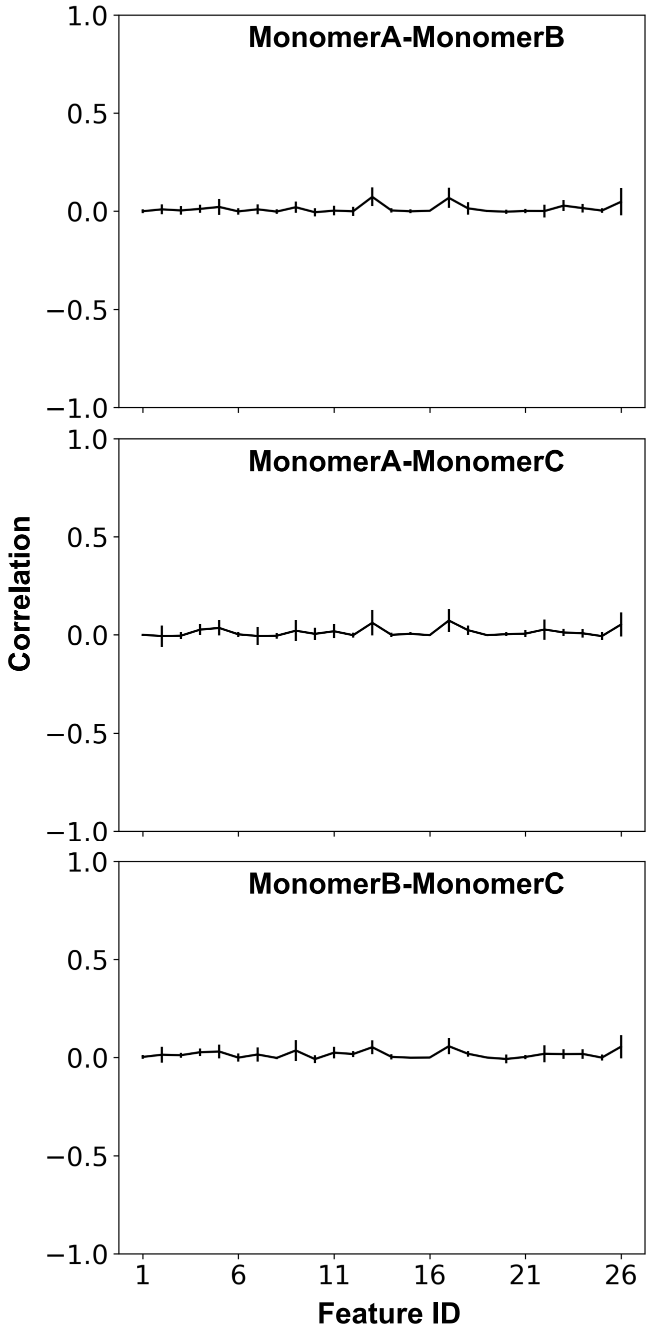


Figure S3: The time-averaged monomer-monomer correlation coefficient for each distance pair feature. The average and standard deviation of the correlation coefficients from all simulations is shown.

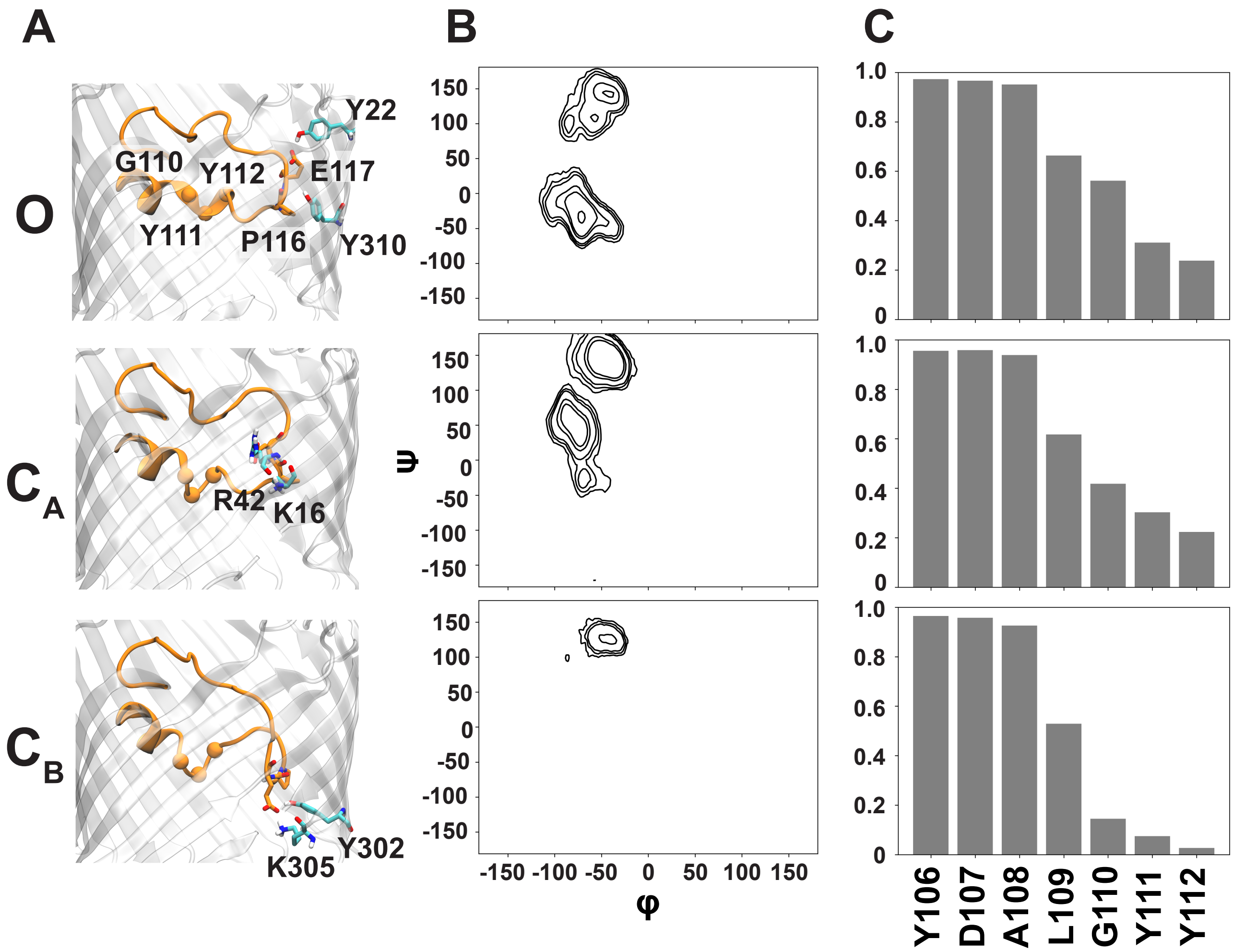


Figure S4: Molecular explanation for the slower $O$-$C_B$ kinetics than $O$-$C_A$ in WT-OmpF. (A) Representative snapshots for $O$, $C_A$ and $C_B$, highlighting the conformation of L3 (colored in orange). The $C_\alpha$ atom for residues belonging to the periplasmic terminal of the helical part of of L3 (G110, Y111, Y112) are shown with VDW representation. (B) Distribution of P116 $\phi/\psi$ angles (in degrees) for the three conformational states, highlighting the conformational restriction of P116 in state $C_B$. (C) The probability of residues located at the periplasmic terminal of L3 in each state to adopt a helical conformation.

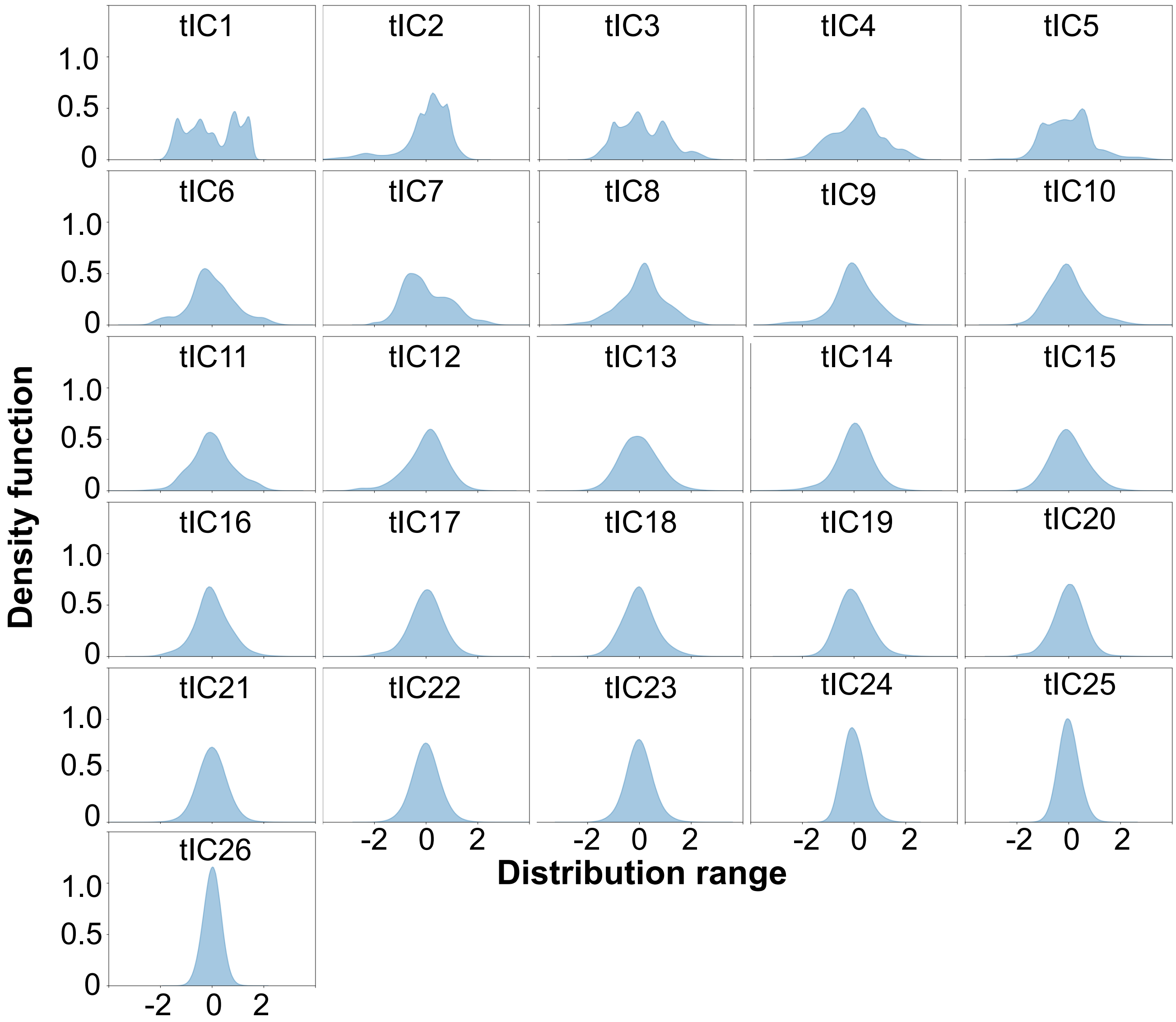


Figure S5: Distributions of all 26 tICA eigenvectors evaluated from all of the distance pair features. The top seven eigenvectors show a non-Gaussian distribution, and are thus sufficient to build the MSM[65]

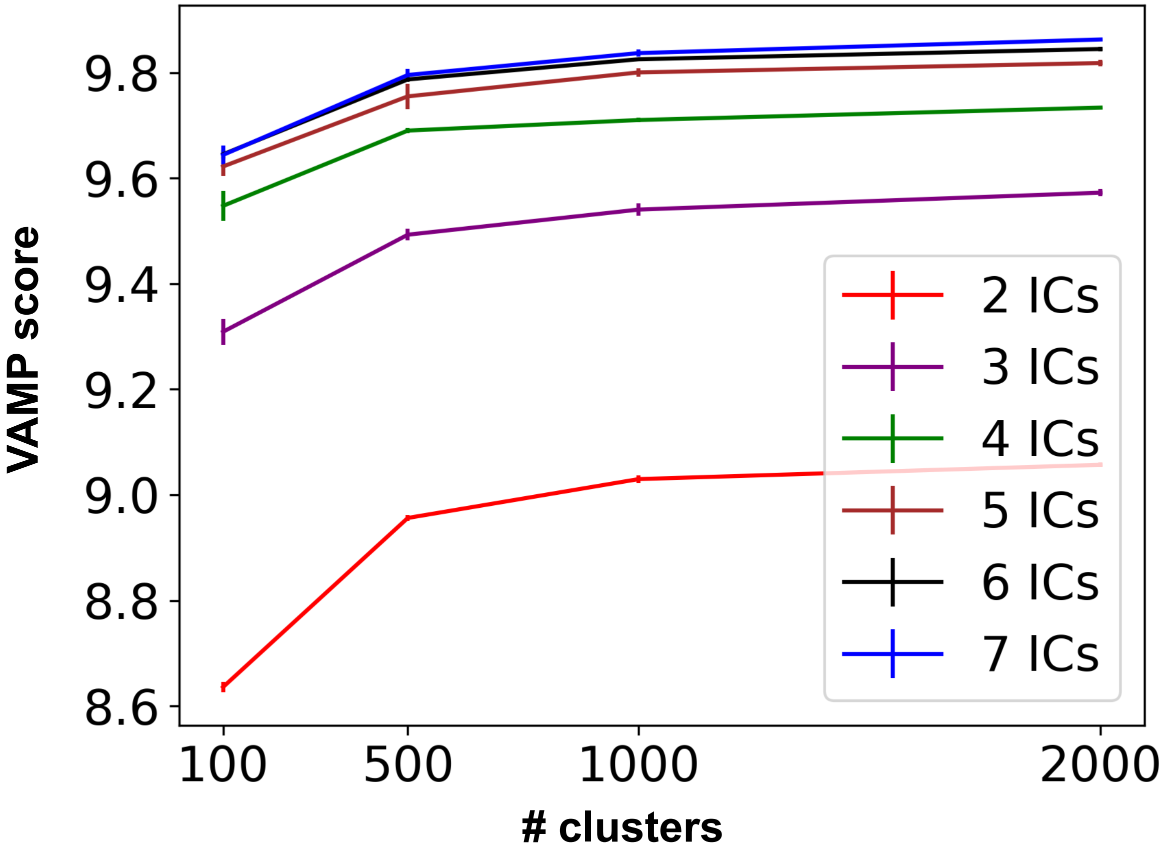


Figure S6: Cross-validated VAMP-2[61] score to rank the parameters (number of tICA eigenvectors and clusters) used to discretize the conformational space. Error bars are calculated by determining the VAMP-2 score after running 5 iterations of the k-means clustering algorithm. The score converges with 5 tICs and 1000 clusters, indicating an optimal parameter set.

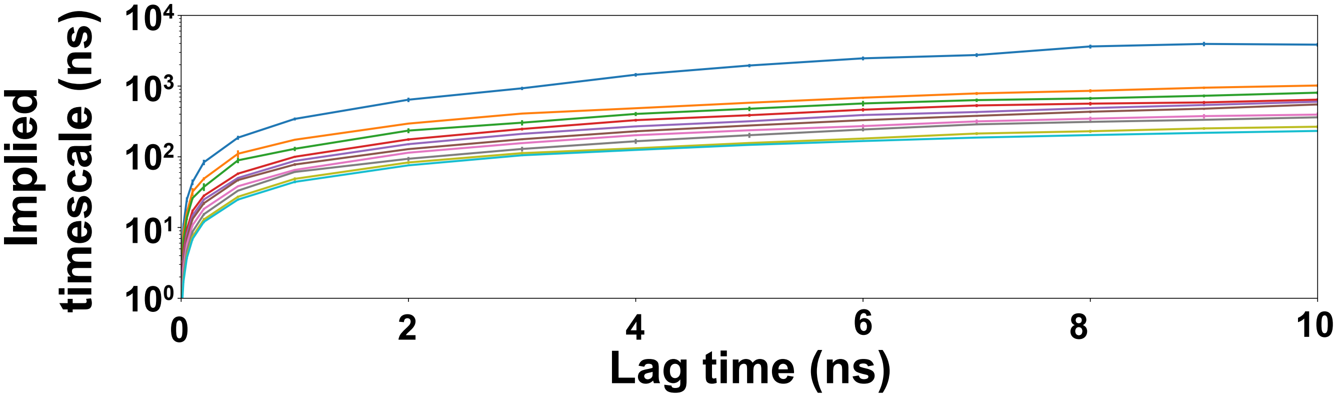


Figure S7: Implied timescales (ITS) plot of the top 10 slowest processes using multiple lag times. Error bars indicate the uncertainty evaluated using a Bayesian estimated MSM. The lag time of 2 ns was chosen for MSM construction as the ITS curve converges, indicating Markovianity.

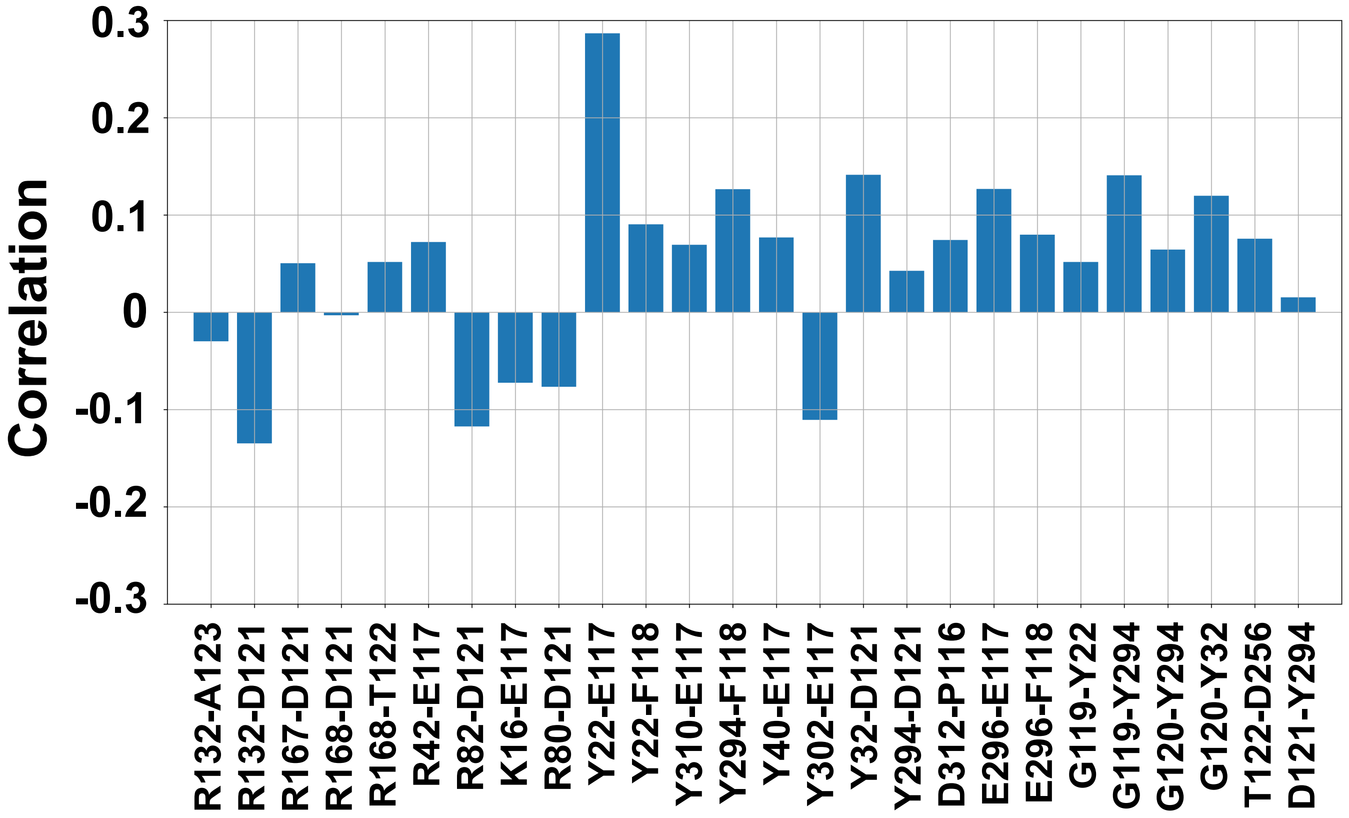


Figure S8: Time-averaged correlation coefficient of each distance pair with the second eigenvector of the TPM to determine optimal indicators of the slowest transition. The Y22-E117 and R132-D121 distance pairs were chosen as indicators since these features had the greatest positive and negative correlation, respectively.

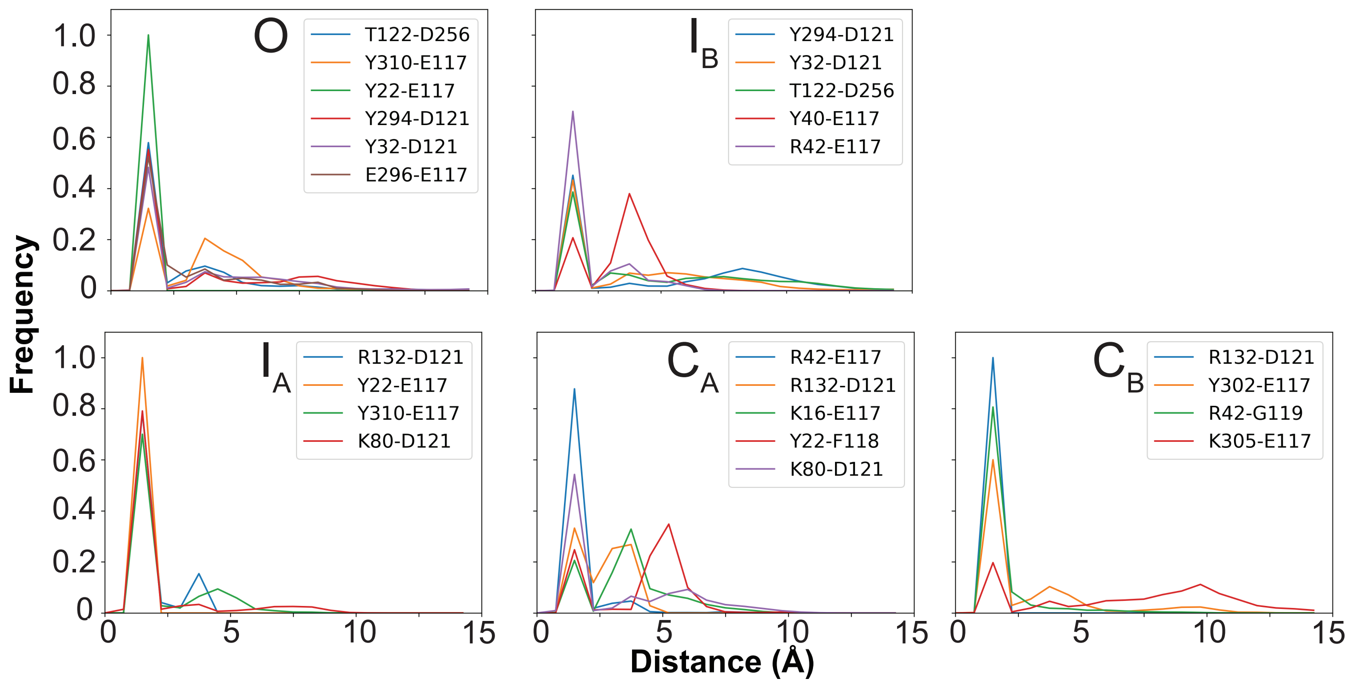


Figure S9: The ensemble distribution of distances between each major hydrogen bond pair (>20 % occupancy) for the 5 conformational states.

```
                       16    22                40 42
  OmpF:  AEIYNKDGNKVDLYGKAVGLHYFSKGNGENSYGGNGDMTYARLGFKGETQINSDLTGYGQ 60
  OmpC:  AEVYNKDGNKLDLYGKVDGLHYFSDNKD-----VDGDQTYMRLGFKGETQVTDQLTGYGQ 55
  PhoE:  AEIYNKDGNKLDVYGKVKAMHYMSDNAS-----KDGDQSYIRFGFKGETQINDQLTGYGR 55
OmpE36:  AEIYNKDGNKLDLYGKAVGLHYFSDNDG-----NDGDKTYARLGFKGETKINDQLTGYGQ 55
OmpK36:  AEIYNKDGNKLDLYGKIDGLHYFSDDKS-----VDGDQTYMRVGVKGETQINDQLTGYGQ 55

                            80                                   116 120
  OmpF:  WEYNFQGNNSEGADAQTGNKTRLAFAGLKYADVGSFDYGRNYGVVYDALGYTDMLPEFGG 120
  OmpC:  WEYQIQGNSAEN-ENN--SWTRVAFAGLKFQDVGSFDYGRNYGVVYDVTSWTDVLPEFGG 112
  PhoE:  WEAEFAGNKAESDTAQ--QKTRLAFAGLKYKDLGSFDYGRNLGALYDVEAWTDMFPEFGG 113
OmpE36:  WEYNFQGNNSEGADAQSGNKTRLAFAGLKFGDAGSFDYGRNYGLVYDAIGITDMLPEFGG 115
OmpK36:  WEYNVQANNTESSSDQ--AWTRLAFAGLKFGDAGSFDYGRNYGVVYDVTSWTDVLPEFGG 113
                                                  |---------------
                                                     L3 residues

        121 122     132
  OmpF:  DTAY-SDDFFVGRVGGVATYRNSNFFGLVDGLNFAVQYLGKNER--------------DT 165
  OmpC:  DTYG-SDNFMQQRGNGFATYRNTDFFGLVDGLNFAVQYQGKNGNPSGEGFTSGVTNNGRD 171
  PhoE:  DSSAQTDNFMTKRASGLATYRNTDFFGVIDGLNLTLQYQGKNEN--------------RD 159
OmpE36:  DTGV-SDNFFSGRTGGLATYRNSGFFGLVDGLNFGVQYLGKNER--------------TD 160
OmpK36:  DTYG-SDNFLQSRANGVATYRNSDFFGLVDGLNFALQYQGKNGSVSGEG----ATNNGRG 168
         --------------|

  OmpF:  ARRSNGDGVGGSISYE-YEG--FGIVGAYGAADRTNLQEAQ-PLGNGKKAEQWATGLKYD 221
  OmpC:  ALRQNGDGVGGSITYD-YEG--FGIGGAISSSKRTDAQNTAAYIGNGDRAETYTGGLKYD 228
  PhoE:  VKKQNGDGFGTSLTYD-FGGSDFAISGAYTNSDRTNEQNLQ-SRGTGKRAEAWATGLKYD 217
OmpE36:  ALRSNGDGWATSLSYD-FDG--FGIVGAYGAADRTNAQQNL-QWGKGDKAEQWATGLKYD 216
OmpK36:  WSKQNGDGFGTSLTYDIWDG--ISAGFAYSHSKRTDEQNSVPALGRGDNAETYTGGLKYD 226

  OmpF:  ANNIYLAANYGETRNATPITNKFTNTSGFANKTQDVLLVAQYQFDFGLRPSIAYTKSKAK 281
  OmpC:  ANNIYLAAQYTQTYNATRVG-----SLGWANKAQNFEAVAQYQFDFGLRPSLAYLQSKGK 283
  PhoE:  ANNIYLATFYSETRKMTPIT------GGFANKTQNFEAVAQYQFDFGLRPSLGYVLSKGK 271
OmpE36:  ANNIYLAALYGEMRNAARLD------NGFANKTQDFSVVAQYQFDFGLRPSIAYYKSKAK 270
OmpK36:  ANNIYLASQYTQTYNATRAG-----SLGFANKAQNFEVVAQYQFDFGLRPSVAYLQSKGK 281

                     294 296   302 305  310
  OmpF:  DVE-GIGDVDLVNYFEVGATYYFNKNMSTYVDYIINQIDSDN---KLGVGSDDTVAVGIV 337
  OmpC:  NLGRGYDDEDILKYVDVGATYYFNKNMSTYVDYKINLLDDNQFTRDAGINTDNIVALGLV 343
  PhoE:  DIE-GIGDEDLVNYIDVGATYYFNKNMSAFVDYKINQLDSDN---KLNINNDDIVAVGMT 327
OmpE36:  DVE-GIGDEDYINYIDIGATYYFNKNMSTYVDYQINQLKDDN---KLGINNDDTVAVGLV 326
OmpK36:  DLERGYGDQDILKYVDVGATYYFNKNMSTYVDYKINLLDDNSFTRNAGISTDDVVALGLV 341

  OmpF:  YQF 340
  OmpC:  YQF 346
  PhoE:  YQF 330
OmpE36:  YQF 329
OmpK36:  YQF 344
```

Figure S10: Sequence alignment of OmpF (*E. coli*), OmpC (*E. coli*), PhoE (*E. coli*), OmpK36 (*K. pneumoniae*) and OmpE36 (*E. cloacae*). Each residue is represented by their one-letter abbreviation. Important residues in the observed gating mechanism of OmpF are highlighted with a box and colored based on residue type (black: hydrophobic, green: polar, red: acidic, and blue: basic). All other residues are colored gray. The alignment was performed on full-length sequences using the Clustal Omega[66] alignment tool within Uniprot.[67]

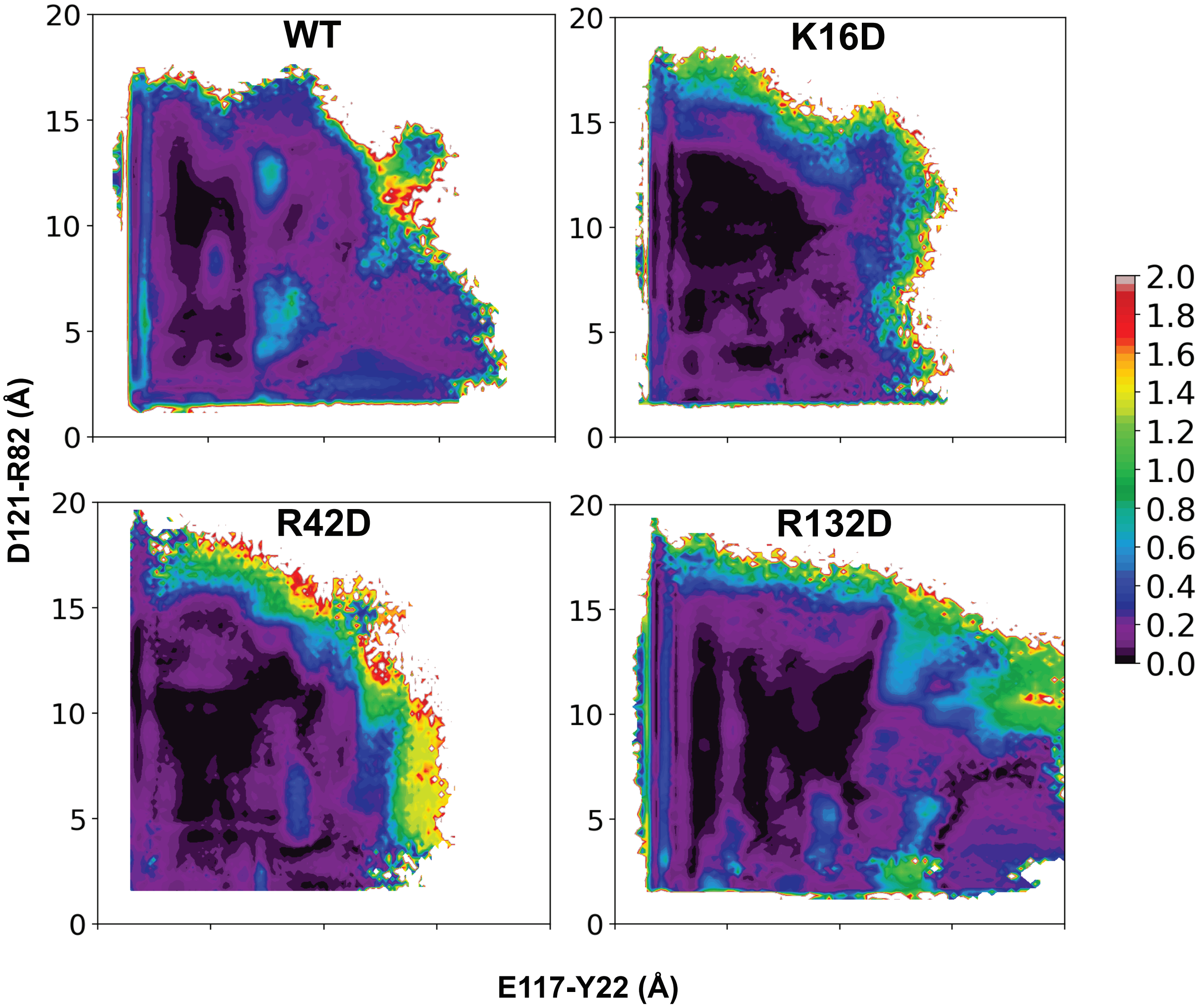


Figure S11: Free energy error for WT, K16D, R42D, and R132D-OmpF systems. The landscape is projected onto E117-Y22 and R132-D121 for WT-OmpF and K16D-OmpF, and E117-Y22 and R82-D121 for R132D-OmpF.

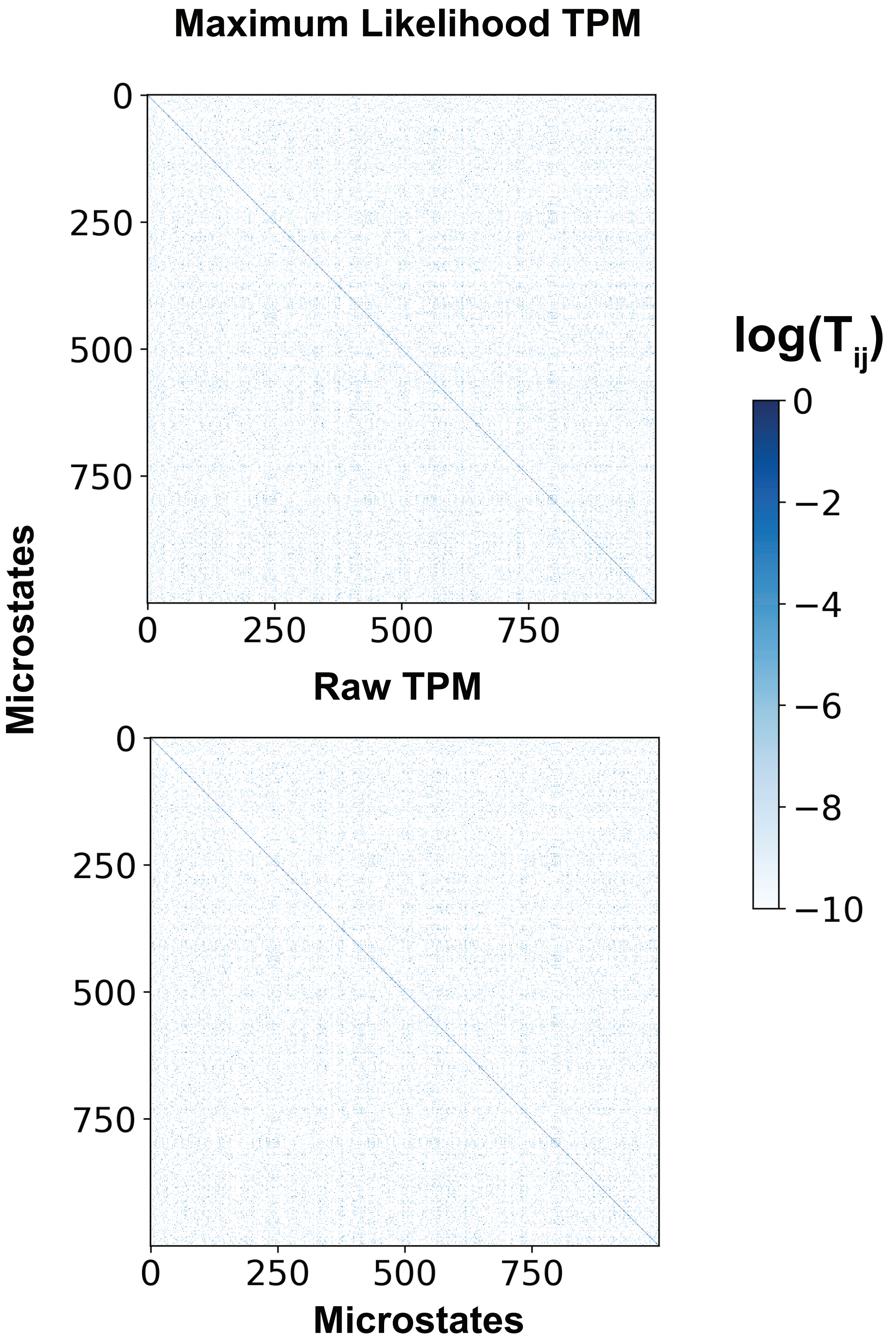


Figure S12: Comparison of the TPM computed with maximum likelihood MSM and with the raw trajectory data. Shown is the log of each matrix element. No significant difference is observed.

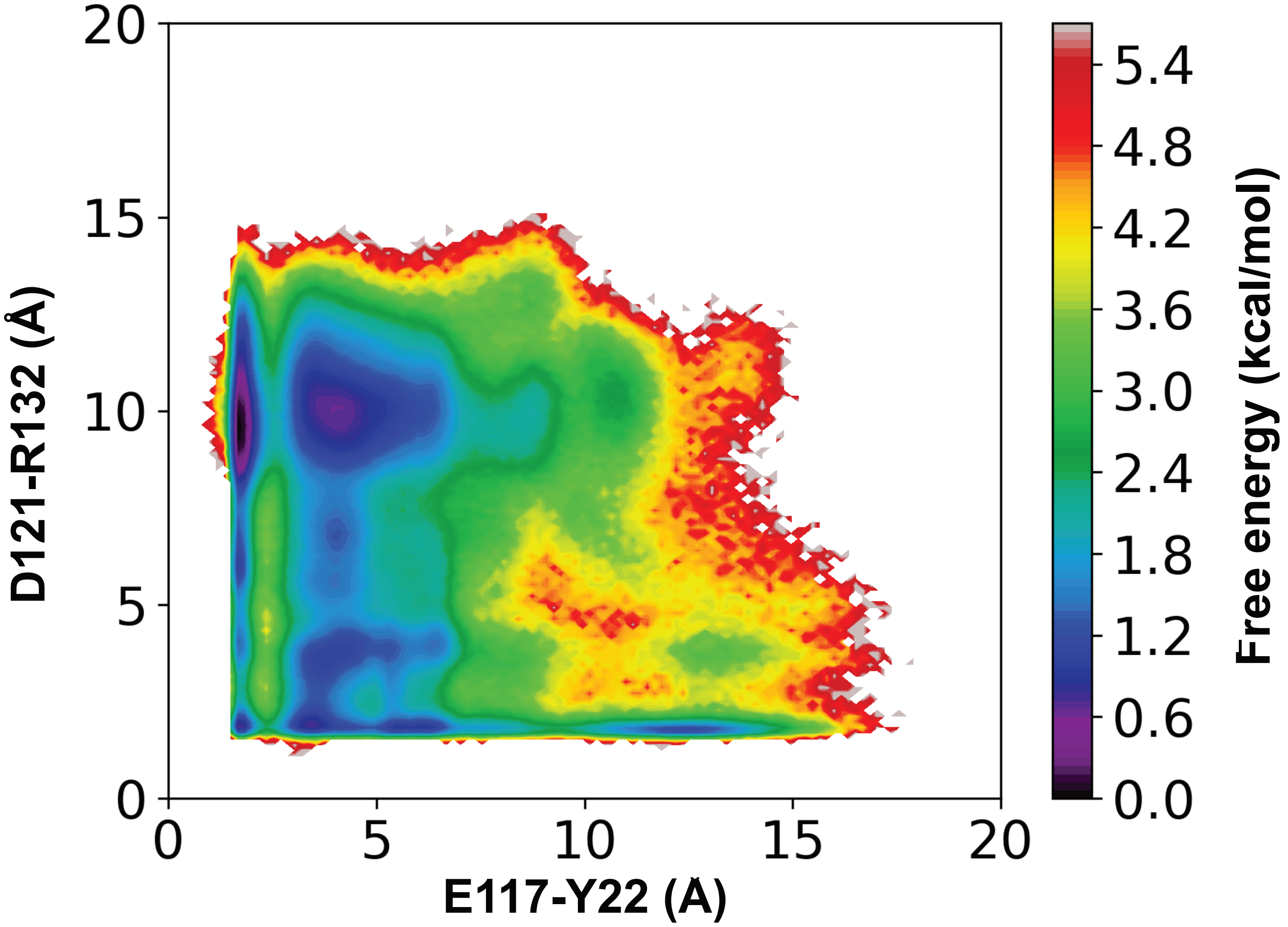


Figure S13: The conformational landscape of L3 projected onto the E117-Y22 and D121-R132 distance features using the raw trajectory data.

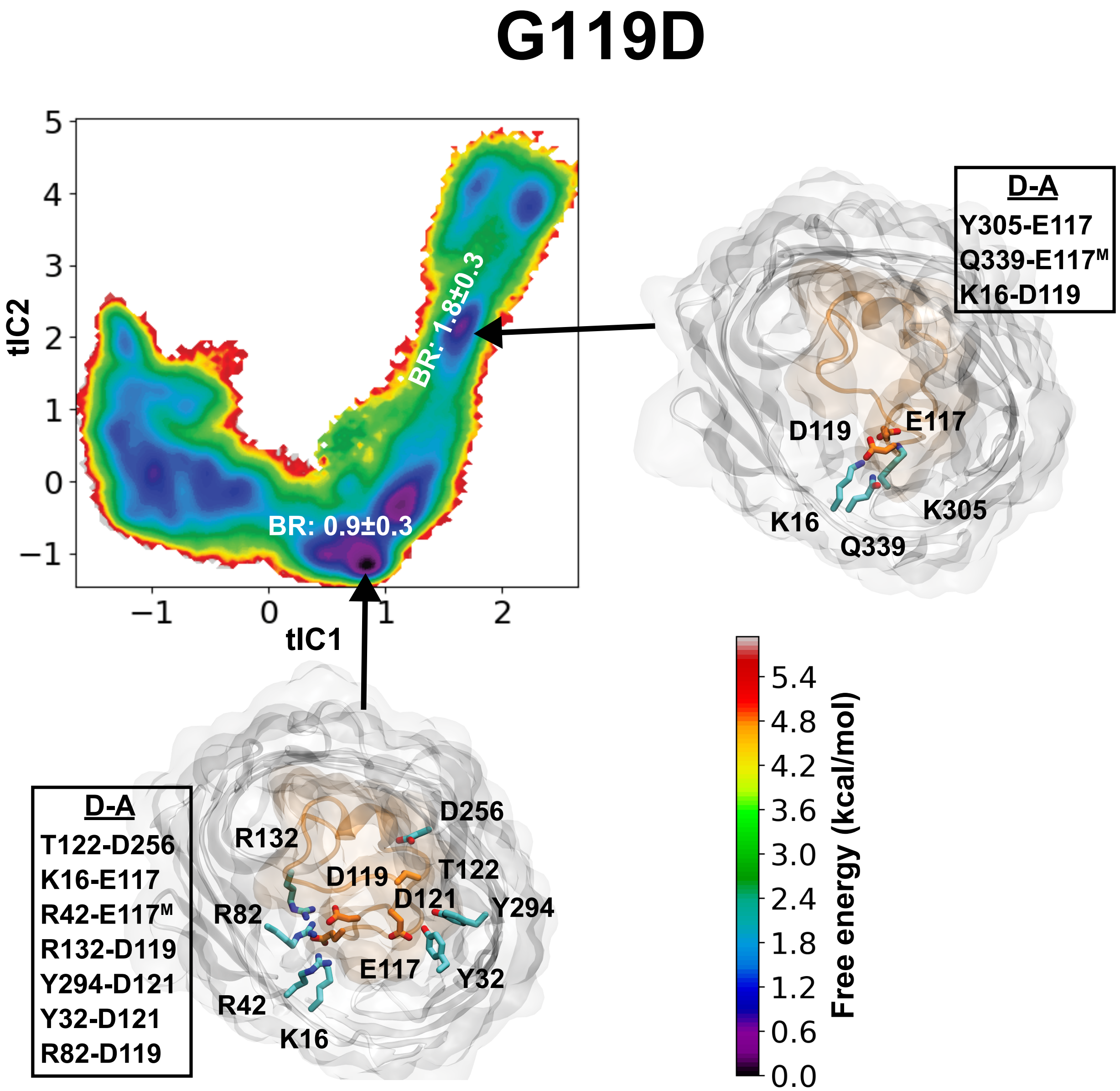


Figure S14: Conformational landscape of L3 in G119D-OmpF mutant. Free energy landscape for dynamics of L3, reweighted by the stationary distribution, is projected onto the top two tICA eigenvectors. The pore bottleneck radii (BR) for the conformational states corresponding to energetic minima are highlighted on the free energy surface. Structural characteristics in each metastable state are depicted by the top-down snapshots of OmpF, highlighting hydrogen bonds with > 20% occurrence probability between the most fluctuating residues of L3 and the barrel residues.

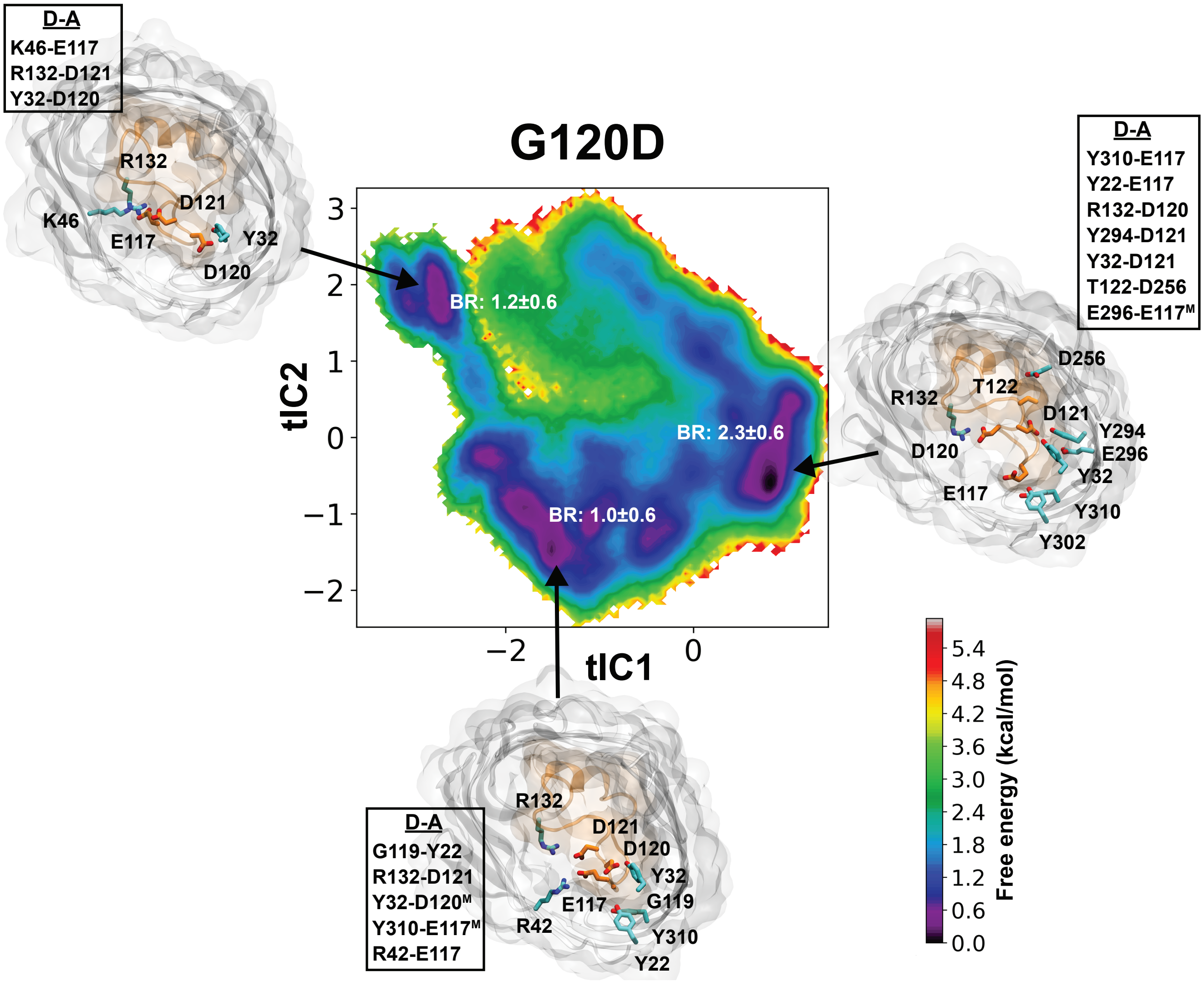


Figure S15: Conformational landscape of L3 in G120D-OmpF mutant. Free energy landscape for dynamics of L3, reweighted by the stationary distribution, is projected onto the top two tICA eigenvectors. The pore bottleneck radii (BR) for the conformational states corresponding to energetic minima are highlighted on the free energy surface. Structural characteristics in each metastable state are depicted by the top-down snapshots of OmpF, highlighting hydrogen bonds with $> 20\%$ occurrence probability between the most fluctuating residues of L3 and the barrel residues.

Table S1: *Escherichia coli* strains used in the study

| **Strain** | **Genotype** |
|---|---|
| BW26678 | *lacI*$^q$ *rrnB*3 Δ*lacZ*4787 *hsdR*514 Δ*araBAD*567 Δ*rhaBAD*568 *rph*-1/pKD46 |
| WM8897 | *lacI*$^q$ *rrnB*3 Δ*lacZ*4787 *hsdR*514 Δ*araBAD*567 Δ*rhaBAD*568 *rph*-1 Δ*ompF*8897::*cat* |
| WM8901 | *lacI*$^q$ *rrnB*3 Δ*lacZ*4787 *hsdR*514 Δ*araBAD*567 Δ*rhaBAD*568 *rph*-1 Δ*ompF*8897::*cat*/pAH69 |
| WM8819 | *lacI*$^q$ *rrnB*3 Δ*lacZ*4787 *hsdR*514 Δ*araBAD*567 Δ*rhaBAD*568 *rph*-1 Δ*ompF*8897::*cat* *att*-HK::pMEM503 |
| WM8820 | *lacI*$^q$ *rrnB*3 Δ*lacZ*4787 *hsdR*514 Δ*araBAD*567 Δ*rhaBAD*568 *rph*-1 Δ*ompF*8897::*cat* *att*-HK::pMEM504 |

Table S2: Plasmids, carrying various *ompF* alleles, used in this study

| **Plasmid** | **Description** | **Construction or Reference** |
|---|---|---|
| pKD3 | Source of *cat*-cassette used for ompF deletion | Datsenko and Wanner [62] |
| pAH144 | *att*-HK integration plasmid, Strep/Spec$^R$ | Haldimann and Wanner[63] |
| pAH69 | Temperature-sensitive helper plasmid for chromosomal integration of *att*-HK plasmids | Haldimann and Wanner[63] |
| pMEM501 | pAH144::*ompF*(wt) | Vector pAH144 cut with EcoRI-HF and SphI-HF (no SAP) ligated with PCR amplified *ompF*(wt) using primers ompF-cloningF and ompFcloningR |
| pMEM503 | pAH144::*ompF*(G120D) | Hi Fi Assembly of MluI/PvuII-cut pMEM501 and ompF-G120D gBlock |
| pMEM504 | pAH144::*ompF*(R132D) | Hi Fi Assembly of MluI/PvuII-cut pMEM501 and ompF-R132D gBlock |

Table S3: Primers used in this study

| **Name** | **Purpose** | **Sequence** |
|---|---|---|
| ompF-cloningF | cloning *ompF*(wt) | GGCGCGCCGCATGCTTCCGTTCC-CACGTACTCCG |
| ompF-cloningR | cloning *ompF*(wt) | GGCGCGCCGAATTCCAGGAGCG-GCGGTAATGTTC |
| del-ompF-F | amplification of *cat*-cassette used for construction of $\Delta$*ompF* mutant | AGATTTTGTGCCAGGTCGATAAA-GTTTCCATCAGAAACAAGTGTAG-GCTGGAGCTGCTTC |
| del-ompF-R | amplification of *cat*-cassette used for construction of $\Delta$*ompF* mutant | GTCCTGTTTTTTCGGCATTTAAC-AAAGAGGTGTGCTATTACATATG-AATATCCTCCTTAG |
| HK022-P1 | verification of single copy plasmid integration at *att*-HK | GGAATCAATGCCTGAGTG |
| HK022-P2 | verification of single copy plasmid integration at *att*-HK | GGCATCAACAGCACATTC |
| HK022-P3 | verification of single copy plasmid integration at *att*-HK. | ACTTAACGGCTGACATGG |
| HK022-P4 | verification of single copy plasmid integration at *att*-HK | ACGAGTATCGAGATGGCA |

Table S4: gBlocks used for construction of *att*-HK plasmids carrying mutated *ompF* alleles

| **Name** | |
|---|---|
| ompF-<br>G120D | CTCTGAAGGCGCTGACGCTCAAACTGGTAACA-<br>AAACGCGTCTGGCATTCGCGGGTCTTAAATACG-<br>CTGACGTTGGTTCTTTCGATTACGGCCGTAACT-<br>ACGGTGTGGTTTATGATGCACTGGGTTACACC-<br>GATATGCTGCCAGAATTTGGTGATGATACTGCA-<br>TACAGCGATGACTTCTTCGTTGGTCGTGTTGGCG-<br>GCGTTGCTACCTATCGTAACTCCAACTTCTTTG-<br>GTCTGGTTGATGGCCTGAACTTCGCTGTTCA-<br>GTACCTGGGTAAAAACGAGCGTGACACTGCACGCC-<br>GTTCTAACGGCGACGGTGTTGGCGGTTCTATC-<br>AGCTACGAATACGAAGGCTTTGGTATCGTTGG-<br>TGCTTATGGTGCAGCTGACCGTACCAACCTGCAAG-<br>AAGCTCAACCTCTT |
| ompF-<br>R132D | CTCTGAAGGCGCTGACGCTCAAACTGGTAACAA-<br>AACGCGTCTGGCATTCGCGGGTCTTAAATACG-<br>CTGACGTTGGTTCTTTCGATTACGGCCGTAACTAC-<br>GGTGTGGTTTATGATGCACTGGGTTACACCGATAT-<br>GCTGCCAGAATTTGGTGGTGATACTGCAT-<br>ACAGCGATGACTTCTTCGTTGGTGATGTTGGCGGCGT-<br>TGCTACCTATCGTAACTCCAACTTCTTTGGTCTGGT-<br>TGATGGCCTGAACTTCGCTGTTCAGT-<br>ACCTGGGTAAAAACGAGCGTGACACTGCACGCCGTT-<br>CTAACGGCGACGGTGTTGGCGGTTCTATCAGCTAC-<br>GAATACGAAGGCTTTGGTATCGTTGGTGCTT-<br>ATGGTGCAGCTGACCGTACCAACCTGCAAGAAGCT-<br>CAACCTCTT |
